# A new class of inherently efficient SUMOylation substrates

**DOI:** 10.64898/2026.09.10.750468

**Authors:** El Hadji Cisse, Laura Visticot, Rafael Cepa, Aanchal Mishra, Franck Coste, Stephane Goffinont, Lucija Mance, Sarah Battault, Dorian Guigneau, Ibtissam Talhaoui, Bertrand Castaing, Vincent Aucagne, Pierre-Antoine Defossez, Marcin Józef Suskiewicz

**Affiliations:** Centre de Biophysique Moléculaire, UPR4301 CNRS, 45071 Orléans Cedex 2, France; Epigenetics & Cell Fate Centre, UMR7216 CNRS, Université Paris Cité, 75013 Paris, France

## Abstract

SUMOylation is an essential eukaryotic ubiquitin-like post-translational modification that plays a central role in the regulation of various nuclear processes and stress responses. It canonically occurs at lysine residues within ΨKXE consensus motifs that lie in intrinsically disordered regions or loops and interact specifically with the SUMO-conjugating E2 enzyme UBC9. However, many detected SUMOylation sites are found within structured domains, and it remains unclear how these are recognised by UBC9. Here, we investigated the SUMOylation of Lys_43_ in the BTB domain of human ZBTB38 (ZBTB38^BTB^), a lysine located within a rigid β-sheet. By combining X-ray crystallography, structural prediction, and *in-vitro* UBC9 interaction and SUMOylation assays, we show that ZBTB38^BTB^possesses a dedicated surface that recapitulates the spatial arrangement of residues found in canonical linear consensus motifs. This surface binds UBC9 with mid-micromolar affinity and is predicted to position Lys_43_ in its active site for efficient SUMOylation. Structural modelling and sequence analyses suggest that this property is shared by BTB domains of five members (10%) of the ZBTB-protein family across vertebrates, revealing a previously unrecognised property of a subset of ZBTB^BTB^domains. Kinetic analyses reveal that, under the reaction conditions used, the catalytic efficiency of ZBTB38^BTB^and ZBTB33^BTB^SUMOylation are closely comparable to that of the C-terminal domain of RANGAP1, the best-characterised and most efficiently SUMOylated substrate known. This defines a new class of inherently efficient, E3 ligase-independent SUMOylation substrates beyond RANGAP1 and suggests that structural pre-organisation of the acceptor lysine and its environment may promote productive UBC9 engagement. Lastly, we demonstrate the presence of higher-molecular-weight, modified forms of ZBTB38 in human cells, consistent with SUMOylation. Together, these results provide a biochemical basis for interpreting existing and designing future studies on the functional impact of ZBTB SUMOylation. More broadly, our findings offer insights into the determinants of efficient SUMOylation, and may facilitate the identification of further inherently efficient targets, and, potentially, the design of SUMOylation modulators.

## Introduction

Post-translational modifications (PTMs) are fundamental regulators of protein function that enable precise spatial and temporal control of cellular processes^1^. Among these, SUMOylation – a PTM related to ubiquitylation – has attracted considerable interest due to its essentiality for organismal viability across eukaryotes^2–5^. At a cellular level, it is involved in diverse fundamental processes, such as genome maintenance, gene expression regulation, nuclear trafficking, and protein degradation^6–10^.

SUMOylation involves a covalent attachment between the side-chain ε-amino group of a lysine residue within a substrate protein and the C-terminal extremity of a small ubiquitin-like modifier (SUMO) proteins^11–13^. The number of SUMO paralogues varies among species, ranging from a single one in *Saccharomyces cerevisiae* (Smt3) to five in humans: SUMO1 to SUMO5, of which SUMO1, SUMO2, and SUMO3 are the most widely expressed and the best characterised^14–17^. Despite notable differences in sequence homology (97% amino-acid identity between SUMO2 and SUMO3, compared with 47% between either of them and SUMO1), all human SUMOs are conjugated through the same core enzymes^18,19^.

The tightly regulated SUMOylation cascade proceeds through four steps: maturation, activation, conjugation, and ligation^11,12^. Initially, SUMO precursors undergo proteolytic maturation mediated by sentrin (a historical name for SUMO^20^)-specific proteases (SENPs) that expose a conserved diglycine motif at the SUMO C-terminus. Mature SUMO then undergoes ATP-dependent activation catalysed by the hetero-dimeric E1 enzyme composed of SAE1 and SAE2 (also known as AOS1 and UBA2, respectively), resulting in a SUMO∼AMP intermediate (where “∼” denotes a reactive covalent bond, in this case a mixed carboxyl-phosphoric anhydride bond). This intermediate subsequently reacts with Cys_173_ of SAE2, resulting in a thioester-linked SUMO∼SAE2 conjugate. The SUMO protein is then transferred from SAE2 onto Cys_93_ of the E2 conjugating enzyme UBC9 (also known as UBE2I), generating a SUMO∼UBC9 intermediate. Lastly, this conjugate directly transfers SUMO onto a lysine side chain within a target protein, yielding SUMO—substrate (where “—” corresponds to a stable iso-peptide bond). SUMOylation is enzymatically reversible and highly dynamic in the cell, with SENPs catalysing deconjugation, thereby carefully regulating the net modification level of substrates.

Generally, E1 and E2 enzymes are sufficient to carry out SUMOylation *in vitro*. However, so far, efficient (within minutes) and quantitative (100%) SUMOylation with E1 and E2 enzymes *in vitro* has only been observed for a single substrate, the C-terminal domain (CTD) of RAN GTPase-activating protein 1 (RANGAP1). The efficient SUMOylation of RANGAP1 enabled the discovery of SUMOylation by several groups studying this protein at the end of the 1990s^21,22^. A few other substrates, including thymine DNA glycosylase (TDG), the ubiquitin-conjugating enzyme UBE2K (also known as E2-25K), and the phosphorylated forms of MEF2 and HSF1 transcription factors, also show above-average SUMOylation efficiency, but are notably less efficiently modified than RANGAP1^23–25^. Most other tested substrates are inefficiently modified with E1 and E2 enzymes *in vitro*, often requiring hours under standard conditions to achieve a low level of SUMOylated product^26,27^.

Although *in-vitro* characterisation is available only for a small number of substrates, proteomics analyses have revealed thousands of proteins with detectable SUMOylation in human cells^28–35^. A recent meta-analysis of existing data identified 35,721 high-confidence SUMOylation sites across 6,146 human proteins^36^, most of them discovered using SUMO2/3-specific protocols.

A part of the answer to the enigma of why most SUMOylation substrates tested to date are inefficiently modified by E1 and E2 enzymes *in vitro* lies in the existence of additional factors in cells, referred to as SUMO E3 ligases^12,37^. *In vitro*, proteins with this activity enhance the transfer of SUMO from the SUMO∼UBC9 conjugate onto lysine residues within substrates. By interacting with the SUMO∼UBC9 conjugate, SUMO E3 ligases act as a structural scaffold, able to stabilise the conjugate in an active conformation, also known as the “closed” conformation^38–41^ – an acceleration mechanism that is analogous to that employed by really interesting new gene (RING)-domain-containing ubiquitin E3 ligases^42–44^. SUMO E3 ligases, which can be induced under specific conditions and localise to particular cellular compartments and substrates, are thought to play a central role in shaping cellular SUMOylation, adding an extra layer of regulation on top of the intrinsic SUMOylability of substrates by the core E1 and E2 enzymes alone^37^.

Large-scale studies analysing thousands of SUMOylation sites revealed that SUMOylation is often found on lysine residues located in intrinsically disordered regions^30,33,34,36^. Besides, SUMOylation tends to occur on a specific linear consensus motif, ΨKXE^45,46^ (where “Ψ” is a hydrophobic residue and “X” is any amino acid residue), or on its variants. Reported motif variants include the form with aspartate instead of glutamate, the inverted consensus orientation^28^, the extended negatively-charged motif (NDSM) containing additional acidic residues at downstream positions, the phosphorylation-dependent SUMOylation motif (PDSM) featuring phosphorylation of a downstream serine or threonine^47–51^, and the motifs containing upstream hydrophobic regions^28,52^. Apart from consensus-motif sites, many non-consensus SUMOylation sites have also been reported, with some of them characterised in more detail and apparently depending on structural conformation rather than linear sequence, as in the case of Lys_14_ of UBE2K^23^ or Lys_1731_ within the SecPH domain of Neurofibromin 1 (NF1)^53^.

A canonical ΨKXE motif is found in RANGAP1 as well, but within a short, conformationally constrained loop protruding from a folded domain rather than a typical intrinsically disordered region. A structural study of the UBC9:RANGAP1^CTD^ complex demonstrated that the consensus motif interacts directly with UBC9^54^. The hydrophobic Ψ residue at the -1-position relative to the modified lysine is recognised through van der Waals contacts by a hydrophobic pocket within UBC9 formed by Pro_128_, Ala_129_, Gln_130_, and Ala_131_. In parallel, the glutamate at the +2-position establishes a hydrogen-bonding and electrostatic-interaction network involving Ser_89_, Thr_91_, Lys_74_, and Lys_76_ at the UBC9 surface. These two interaction sites position the target lysine residue close to the catalytic Cys_93_ residue carrying thioester-linked SUMO, thereby enhancing SUMO ligation. In addition to the consensus motif, RANGAP1^CTD^ interacts with UBC9 *via* an additional surface, contributing to the elevated affinity and modification of this model substrate.

Several studies, including both high-throughput proteomics and dedicated studies, have identified ZBTB proteins as SUMOylation substrates in cells^55,28,56,30,29,31,57,58,33,59–61^, although, to our knowledge, their SUMOylation has not been studied *in vitro* to date. The ZBTB protein family, counting around fifty distinct proteins in humans, constitutes an evolutionarily conserved class of transcriptional repressors implicated in cell differentiation and development^62–64^ and, in a disease context, in cancer^63,65,66^ and developmental syndromes^67,68^. As their name suggests, ZBTB proteins contain two distinct sets of domains, zinc finger (ZF) and broad-complex, tram-track, and bric-à-brac (BTB) domains, connected by a linker region. The multiple C-terminal ZF domains mediate DNA binding and determine specificity toward particular target DNA sequences. By contrast, the BTB domain – usually present as a single copy at the N-terminus – binds co-repressors and promotes multimerisation, ranging from a simple homo- or hetero-dimer formation to higher-order assemblies^69–75^.

Recently, our group and another team have independently characterised a subset of ZBTB proteins – estimated to include around twenty distinct proteins (40% of the ZBTB family) in humans – whose BTB domains form filamentous structures composed of dimers^71,73^. Motivated by this discovery, we aimed to structurally characterise the BTB domains of those members of the ZBTB family for which structural information remains limited. Here, we focused on the BTB domain of ZBTB38 (ZBTB38^BTB^), a methyl-DNA-binding ZBTB protein^76^. We determined the crystal structure of ZBTB38^BTB^ and demonstrated that, while it does not detectably form higher-order assemblies, it contains an evolutionarily conserved surface that directly recruits the SUMO-conjugating E2 enzyme UBC9. The identified surface contains both an acidic residue and a few hydrophobic ones, mimicking, in three dimensions, a linear consensus SUMOylation site. This transient interaction, reconstituted and quantified *in vitro*, results in SUMOylation of the conserved Lys_43_ residue, a site previously identified by high-throughput proteomics in cells^28,33,77^ but not investigated further. Surprisingly, ZBTB38^BTB^ SUMOylation is as efficient as that of RANGAP1^CTD^ under the applied reaction conditions involving only E1 and E2 enzymes *in vitro*. An equivalent surface is present in around 10% of the ZBTB-family members, including also ZBTB4, ZBTB14, ZBTB21, and ZBTB33 (also known as KAISO), across vertebrates. As the functional relevance of BTB SUMOylation in ZBTB33 and ZBTB14 has previously been demonstrated in human cells or in fish^59,61^, our study provides a plausible biochemical basis for those past functional findings, in addition to exploring ZBTB38 SUMOylation for the first time. Combining X-ray crystallography, structural modelling, and biochemical and cellular assays, our work reveals unprecedentedly efficient SUMOylation of a class of non-RANGAP1 substrates, with potential biological consequences for transcriptional regulation.

## Results

### Crystal structure of ZBTB38BTB dimer

We began this study by recombinantly producing a fragment of human ZBTB38^BTB^ corresponding to residues 1-134 fused to a non-cleavable C-terminal His-tag in *Escherichia coli*. The expected molecular weight (MW) of a single ZBTB38^BTB^ protomer is 16 kDa. During the last purification step, preparative size-exclusion chromatography (SEC), the protein eluted at an apparent MW of 35 kDa, according to column calibration, indicating an expected tendency to homo-dimerise. Of note, BTB domains of ZBTB proteins generally homo-dimerise in a co-translational manner^78^, and the dimeric form has hallmarks of an obligate dimer^69^.

Following purification, the crystallisation of the protein was performed. The crystal structure of ZBTB38^BTB^ was determined at a resolution of 1.95 Å in the C222_1_ space group (**Table 1**). The asymmetric unit contains a single protomer with residues 3-123 resolved in electron density. The core of the protomer comprises three β-strands (β_2_-β_4_) and five α-helices (α_2_-α_6_) folded into an architecture typical of BTB domains and related potassium channel tetramerisation domains (KCTD domains), with a three-stranded β-sheet capped on one side by several α-helices^69,74^ (**Figure 1A**). Notably, ZBTB38^BTB^ also features an N-terminal extension characteristic of ZBTB-type BTB domains comprising the α_1_ helix. While this extension seemingly protrudes into solution in an isolated protomer, analysis of the crystal lattice demonstrates that this region engages a symmetry mate, contributing to the formation of a C2-symmetric homo-dimer similar to those previously observed for other BTB domains of ZBTB and related proteins (**Figure 1B**). First, an intermolecular antiparallel β-sheet is formed through the association of the very N-terminus of one protomer (the β_1_ strand) with a region between the α_4_ and α_5_ helices of the neighbour (the β_5_ strand). Second, dimer formation is reinforced by a network of hydrophobic contacts involving the α_1_ helix of one protomer, which packs against the α_1_ helix and the core fold of the partner. Together, these contacts – burying 1805 Å^2^ of interface surface, featuring twenty-five hydrogen bonds, and showing the estimated solvation free energy gain of -22.5 kcal/mol – ensure the proper assembly and structural stability of the ZBTB38^BTB^ homo-dimer.

**Table 1:** X-ray data collection and refinement statistics.

| **Data collection statistics** |  |
| --- | --- |
| Radiation source | ID23-2 |
| Wavelength (Å) | 0.87313 |
| Spacegroup | C222<sub>1</sub> |
| cell dimensions: a, b, c (Å) | 37.53, 130.14, 50.79 |
| Number of proteins / asymmetric unit | 1 |
| Resolution range (Å) | 36.06-1.95 (2.00-1.95) |
| Total observations | 61814 (4454) |
| Unique reflections | 9463 (661) |
| Completeness (%) | 99.9 (100.0) |
| Multiplicity | 6.5 (6.7) |
| $R_{\text{merge}}$ (%) | 8.1 (140.6) |
| $R_{\text{pim}}$ (%) | 3.5 (57.9) |
| $I/\sigma(I)$ | 11.8 (1.3) |
| $CC_{1/2}$ (%) | 99.9 (60.0) |
| **Refinement statistics** |  |
| Resolution range (Å) | 36.06-1.95 |
| Number of reflections used | 9441 |
| $R_{\text{work}} / R_{\text{free}}$ (%) | 21.52/23.77 |
| Average B values (Å<sup>2</sup>) |  |
| All atoms | 51.10 |
| Protein chain atoms | 51.30 |
| Water atoms | 44.40 |
| Root mean square deviation from ideality |  |
| Bond lengths (Å) | 0.007 |
| Bond angles (°) | 0.777 |
| Ramachandran analysis |  |
| Favoured regions / Allowed regions / Outliers (% of total) | 98.3/1.7/0.0 |
| Number of atoms |  |
| Protein chain | 907 |
| Water | 27 |
| **PDB code** | **32MN** |

**Figure 1.**
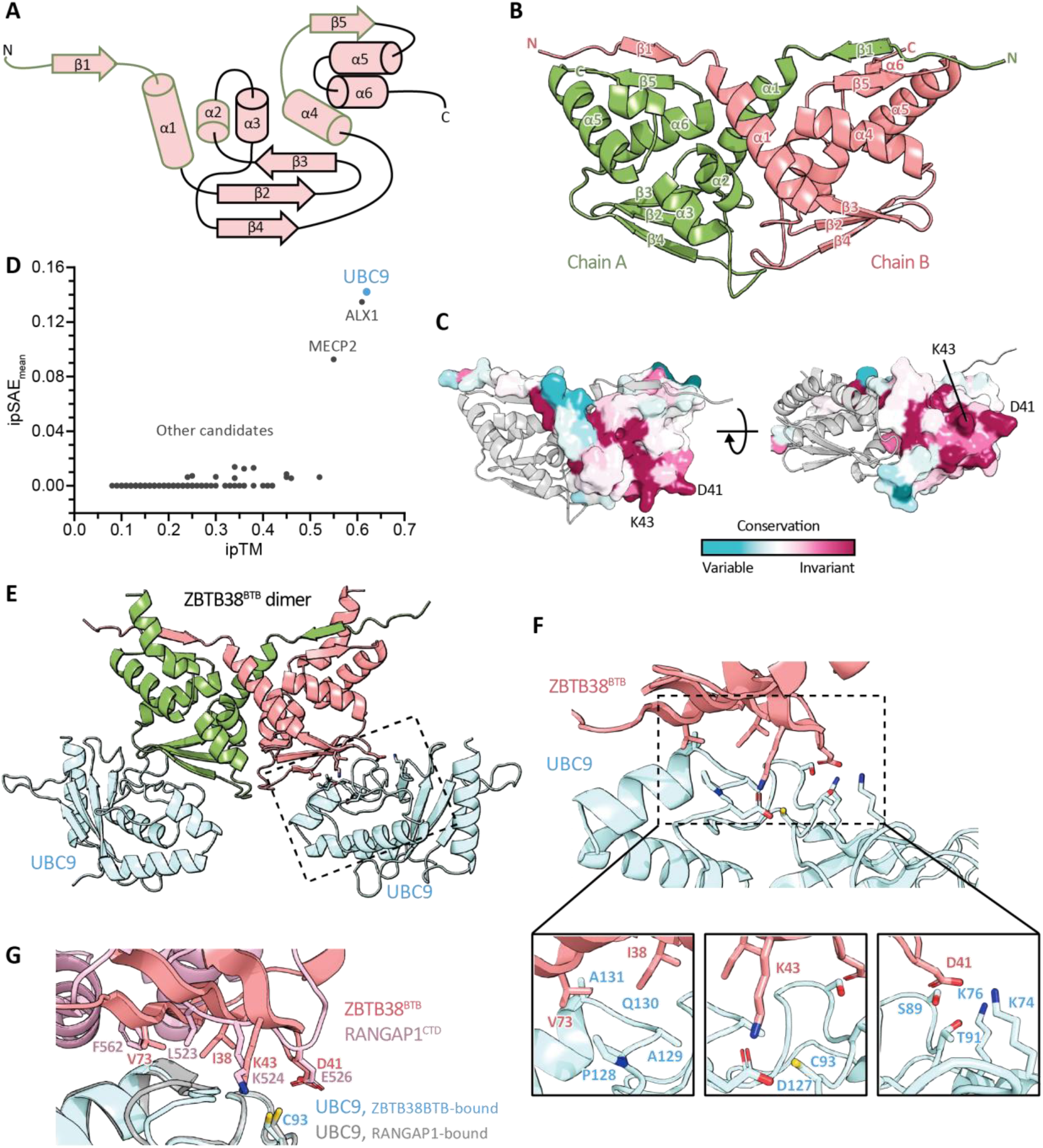
Structural characterisation of the ZBTB38^BTB^ dimer and AlphaFold3 prediction of its interaction with SUMO E2 enzyme UBC9. (A) Topology diagram of ZBTB^BTB^ domains in an orientation corresponding to chain B in panel B. The green frame highlights regions involved in intra-dimer contacts. (B) X-ray diffraction crystal structure of the human ZBTB38^BTB^ dimer, resolved at 1.95 Å in C222_1_ space group and shown as cartoon. The asymmetric unit comprises a single chain, and the dimer is formed from two units *via* C2 crystallographic symmetry. (C) Sequence conservation mapped onto the surface of chain B of ZBTB38^BTB^ according to the indicated colour scale, with chain A shown as grey cartoon. (D) Scatter plot representing the results of computational screening of candidate partners against the ZBTB38^BTB^ dimer, plotted according to ipTM and ipSAE_mean_ predicted confidence metrics. (E) AlphaFold3 model of the ZBTB38^BTB^:UBC9 complex in 2:2 stoichiometry. A part zoomed-in in panel F is marked with a dashed box. (F) Close-up views of the ZBTB38^BTB^:UBC9 interface including interface overview (top), van der Waals contact (bottom left), the target Lys_43_ residue (bottom middle), and polar/electrostatic contact (bottom right). See Supplementary Figure 3 for the predicted local distance difference test (pLDDT) scores and the predicted aligned error (PAE) plot. (G) Structural alignment of the AlphaFold3-predicted ZBTB38^BTB^:UBC9 model with the experimentally determined RANGAP1^CTD^:UBC9 complex (PDB 5D2M), super-imposed on UBC9 molecules.

We further analysed the crystal lattice in search of potential biologically relevant higher-order contacts between dimers. The lattice does not display the inter-dimer contacts mediated by α5 and α6 helices that we and others have shown to drive the higher-order filamentous assemblies in solution for a subset of ZBTB^BTB^ domains^71,73^. It also lacks the extended β1 strand-mediated contacts seen for the BTB domains of ZBTB27/BCL6,^79^ ZBTB16/PLZF^80,81^, ZBTB7A/LRF^82^, which have been speculated to reflect alternative physiological higher-order assemblies, but have not been validated in solution. In fact, the second most extended contact in the ZBTB38^BTB^ lattice after homo-dimerisation does involve β_1_ strands of protomers from two neighbouring dimers but buries only 213 Å^2^ interface area and is unlikely to be biologically relevant, except perhaps where the protein is very highly concentrated locally.

To further explore the homo-oligomerisation propensity of ZBTB38^BTB^, AlphaFold3^83,84^ was used to model a dimer as well as a higher-order tetrameric state. The predicted dimer model has high confidence, as indicated by the interface predicted template modelling (ipTM) score of 0.83 for a top of five models and a predicted aligned error (PAE) plot with dark off-diagonal squares, and is very similar to the experimental structure we resolved, super-imposing with it almost perfectly (root-mean-square-deviation of 0.6 Å over 221 core C_α_ atoms, **Supplementary Figure 1A**). High-confidence dimer prediction was also apparent in the tetramer model; however, the lowered global ipTM score of 0.32 for the top model and the PAE plot with light off-diagonal squares indicate that the predicted interaction between dimers is of low confidence (**Supplementary Figure 1B**). Taken together, the lack of extended crystal contacts beyond homo-dimerisation and the AlphaFold3 modelling results suggest that ZBTB38^BTB^ exists mainly as a dimer and may be unable to form higher-order structures to a meaningful extent. Consistent with these observations, we see no sign of ZBTB38^BTB^ assemblies larger than dimer in preparative SEC (as mentioned above) or in analytical SEC (see below).

### ZBTB38^BTB^ and related BTB domains exhibit a conserved potential interaction surface

To further investigate the structural basis of ZBTB38^BTB^ function, we examined evolutionary conservation across vertebrate orthologues to identify conserved surfaces that might indicate functional interaction sites. The overall high degree of sequence conservation across the whole ZBTB38^BTB^-domain sequence, reflecting strong evolutionary constraint, complicated the discrimination of residues under exceptional constraint indicative of critical functional properties. Therefore, we extended our analysis to BTB domains from ZBTB4, ZBTB14, ZBTB21, and ZBTB33 – ZBTB proteins that belong to the same phylogenetic clade^73^. While each of these homologues is also individually highly conserved across vertebrates, the overall homology across the BTB domain within the clade is moderate (∼40% amino-acid identity), providing sufficient sequence diversity to resolve particularly conserved surfaces that might reflect critical interaction sites shared among all clade members. Generating a multiple sequence alignment of these five ZBTB proteins across vertebrates and quantifying conservation using ConSurf^85^, we mapped BTB-domain regions under strong evolutionary constraint within this clade onto the ZBTB38^BTB^ dimer structure (**Figure 1C**). This analysis revealed a surface corresponding to the β2-β3-β4 β-sheet, located at the bottom of our representation, as particularly conserved. Interestingly, this surface features a lysine residue (Lys_43_ in ZBTB38) that has been previously reported to be SUMOylated in all five clade members^29–31,33,57–61^.

### AlphaFold3 screen suggests that the conserved ZBTB38^BTB^ surface recruits SUMO E2 UBC9

To explore the function of the identified conserved surface, we performed a computational protein:protein interaction screen using a custom framework based on AlphaFold3. Given that BTB domains form tight, likely obligate dimers, we set out to perform the screen with the dimeric ZBTB38^BTB^ form as bait. To simplify the modelling task, we first verified that the dimer could be effectively predicted by AlphaFold3 from a single sequence composed of two BTB-domain sequences repeated in tandem. This approach was inspired by a previous crystallographic study in which fusing two BTB domains in tandem reconstituted well-folded dimers *in vitro*, consistent with the N-terminus of one subunit being close to the C-terminus of the other^86^. Using this single-chain ZBTB38^BTB^ dimer as a structural bait in our screen meant that each individual interaction prediction would be a binary one between the bait and a candidate protein, with interface confidence metrics reflecting the contact between the two unaffected by the BTB intra-dimer interaction. With this approach, we screened the ZBTB38^BTB^ dimer against 153 reported interactors of full-length ZBTB38 listed in BioGRID, a publicly available database aggregating the results of published high- and low-throughput interaction studies^87^. The majority of the tested candidates came from our previous interactomics analysis of ZBTB38^88^. Additionally, motivated by the location of the SUMOylated Lys_43_ residue within this region, we included the SUMO E2 enzyme UBC9 among the interaction candidates, despite UBC9 not having been reported as a ZBTB38 interactor to date. The results were ranked according to the ipTM score and a recently proposed correction to this metric, the interaction prediction score from aligned errors (ipSAE)^89^ (**Figure 1D**). As the ipSAE score is non-commutative (values for the A:B interaction differ depending on whether A or B is considered as the immobile reference), we computer the mean value, ipSAE_mean_.

We selected three top-scoring candidate proteins for visual inspection of their ZBTB38^BTB^-bound models (**Supplementary Figure 2A**). Among these, only UBC9 – the highest-scoring hit (ipTM 0.62, ipSAE_mean_ 0.142) – was predicted to interact with the conserved bottom surface of ZBTB38^BTB^. Strikingly, in the AlphaFold3 model, Lys_43_ was placed within the active site of UBC9, in direct proximity of its catalytic Cys_93_, as expected for a target lysine undergoing SUMOylation. These features, together with the moderate but reproducible confidence scores and the conserved character of the engaged ZBTB38 surface, strongly suggest that the ZBTB38^BTB^:UBC9 model represents a real-world interaction. The other top-scoring interactors, including ALX1 (ipTM 0.61, ipSAE_mean_ 0.135) and MECP2 (ipTM 0.55, ipSAE_mean_ 0.093), contact other ZBTB38^BTB^ regions in the models and their interaction modes are, in our view, less structurally convincing. Here, we focussed on the predicted interaction with UBC9, leaving others for future experimental testing. To test the robustness of the predicted ZBTB38^BTB^:UBC9 pose, we repeated this specific modelling ten times with different seeds. This led to essentially the same model each time and ipTM and ipSAE_mean_ scores within relatively narrow ranges, suggesting robust prediction. (**Supplementary Figure 2B**).

In our AlphaFold3 screen the ZBTB38^BTB^ domain was provided as a single-chain dimer – and modelled against single copies of each candidate protein including UBC9, thus producing a dimeric ZBTB38^BTB^:monomeric UBC9 model that might not represent the real partner stoichiometry. Therefore, we asked whether, structurally, each protomer in the dimer could individually bind one UBC9 molecule without steric clashes. Indeed, modelling of a ZBTB38^BTB^ dimer with two UBC9 molecules using AlphaFold3 revealed binding of two UBC9 molecules, each contacting one BTB-domain copy (**Figure 1E** and **Supplementary Figure 3**). Given the lack of structural constraints, we expect a 2:2 interaction between ZBTB38^BTB^ and UBC9 to be the most likely.

By closely examining the interface of the predicted ZBTB38^BTB^:UBC9 complex (**Figure 1F, *top*** and **Supplementary Figure 3**)., we noticed that the bottom ZBTB38^BTB^ surface mediating UBC9 recognition appears to structurally mimic a SUMO consensus motif. However, whereas SUMO consensus motif residues are contiguous in the primary structure, the equivalent residues in ZBTB38^BTB^ are brought into proximity through the three-dimensional folding of ZBTB38^BTB^. Two sub-sites flanking the target lysine appear to be involved in UBC9 recruitment. First, the hydrophobic Ile_38_ and Val_73_ residues of ZBTB38^BTB^ establish van der Waals contacts with the hydrophobic pocket of UBC9 around Pro_128_ and Ala_129_ (**Figure 1F, *bottom left***). Second, hydrogen bonds and electrostatic interactions connect Asp_41_ of ZBTB38^BTB^ with Ser_89_, Thr_91_, Lys_74_, and Lys_76_ of UBC9 (**Figure 1F, *bottom right***). Moreover, as mentioned above, Lys_43_ is embedded in the catalytic site of UBC9 in a striking way (**Figure 1F, *bottom middle***). The ε-amino group of Lys_43_ is well-positioned both to carry out a nucleophilic attack on the thioester formed at Cys_93_ of UBC9 (distance 2.8 Å) and to interact with Asp_127_ of UBC9, a residue previously shown to enhance the nucleophilicity of the acceptor lysine^26,90^. The Lys_43_:Asp_127_ interaction is likely to also contribute to the overall substrate:UBC9 affinity.

Aligning, on UBC9, the predicted ZBTB38^BTB^:UBC9 complex with the RANGAP1^CTD^:UBC9 complex resolved by X-ray crystallography shows that ε-amino group of both target lysines align very closely, as do carboxyl groups of Asp_41_ of ZBTB38^BTB^ and Glu_526_ (the +2 glutamate of the ΨKXE motif) of RANGAP1^CTD^ (**Figure 1G**). Furthermore, Ile_38_ and Val_73_ are in similar positions to those of Leu_523_ (the –1 hydrophobic residue of the ΨKXE motif) and the auxiliary hydrophobic Phe_562_ of RANGAP1^CTD^. This confirms the structural mimicry of the linear SUMOylation motif by the structural motif of ZBTB38^BTB^.

Lastly, we scrutinised the pre-calculated full-length ZBTB38 model deposited in the AlphaFold Protein Structure Database^91^ to evaluate whether the UBC9-interacting surface is predicted to be accessible in the full-length context. In this model, the BTB domain is directly followed by a long, predicted intrinsically disordered region that is unlikely to occlude the site, and no high-confidence intramolecular contacts between the BTB domain and any other part of the protein that might do so are detectable. In summary, structural modelling robustly predicts an interaction between a ZBTB38^BTB^ dimer and two UBC9 molecules, which should be available in the full-length ZBTB38 context.

### Characterisation of the ZBTB38BTB:UBC9 interaction dependent on the conserved BTB surface

To validate the predicted interaction, we performed analytical SEC experiments. The purified ZBTB38^BTB^ and UBC9 proteins were initially injected independently at 250 µM (here and throughout the article the stated concentrations represent monomer concentration) onto a Superdex75 Increase column at 100 mM NaCl (**Figure 2A-C, *top***). In addition to monitoring ultraviolet (UV) absorbance, the eluate was collected in 1-mL fractions and analysed by sodium dodecyl sulphate polyacrylamide gel electrophoresis (SDS-PAGE) (**Figure 2A-C, *bottom***). ZBTB38^BTB^ eluted as a single peak at 10.90 mL, consistent with a dimer according to column calibration. This confirms the stable dimeric form of ZBTB38^BTB^ while arguing against the formation of higher order multimers, at least at the concentration studied. UBC9 also eluted predominantly in one peak, centred at 12.80 mL and consistent with a monomer.

**Figure 2.**
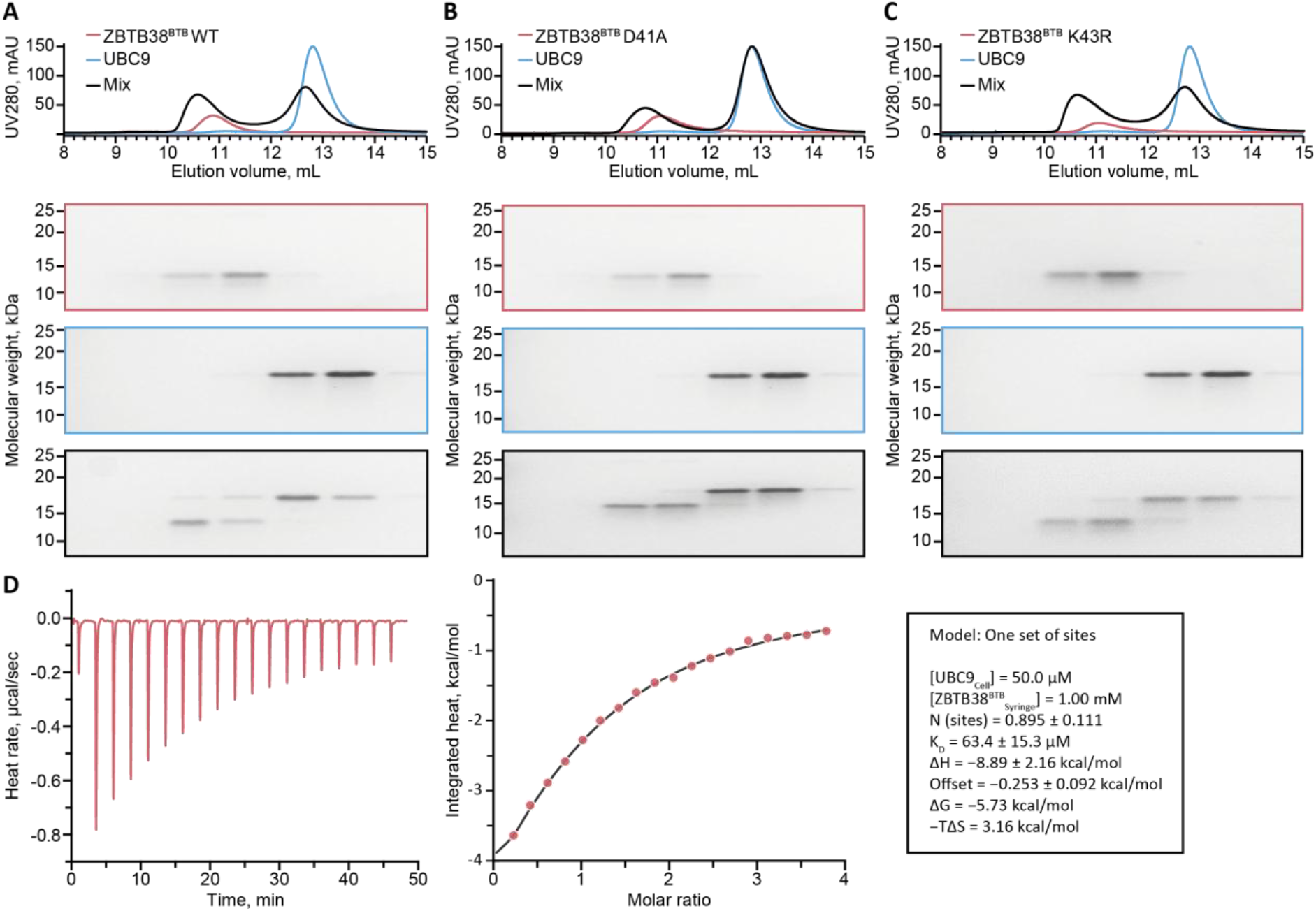
Characterisation of the ZBTB38^BTB^:UBC9 interaction. (A-C) Superposed SEC elution profiles (top) and corresponding fraction analysis by SDS-PAGE (bottom) of the indicated protein samples injected separately or as a mixture. UV280 indicates absorbance at 280 nm. (D) ITC binding isotherm for the titration of ZBTB38^BTB^ into UBC9 (left) and the corresponding integrated enthalpy change per mole of ZBTB38^BTB^ as a function of the molar ratio [ZBTB38^BTB^]:[UBC9] (middle). Data fitting results are provided in the box (right).

The analysis of the protein mixture, with each protein at 250 µM and preincubated at room temperature for 20 min, produced a chromatogram displaying two peaks rather than a single species that would be expected for stable complex formation (**Figure 2A, *top***). However, signs of a transient interaction were readily evident. While the first peak can be predominantly attributed to ZBTB38^BTB^ and the second to UBC9, a shift in elution volume was observed for both peaks (10.57 and 12.67 mL), with a shallower valley between the two peaks that could indicate an equilibrium between the bound and dissociated forms. Moreover, the UV intensity of the first peak increased whereas that of the second decreased, which – based on the SDS-PAGE analysis – can be attributed to the partial earlier elution of UBC9 together with ZBTB38^BTB^ (**Figure 2A, *bottom***).

To quantify this interaction, we performed isothermal titration calorimetry (ITC). The binding isotherm revealed a specific interaction between ZBTB38^BTB^ and UBC9, which could be fitted to a One Set of Sites binding model with estimated stoichiometry close to one and a dissociation constant (*K*_D_) of approximately 60 µM (**Figure 2D**). We observed no obvious signs of cooperativity, but cooperativity was not formally quantified, and we assumed a non-cooperative model for fitting. Given the low *c*-value of this ITC measurement, the fitted stoichiometry and *K*_D_ should be considered approximate and model-dependent; nonetheless, the obtained stoichiometry is consistent with the AlphaFold3 model, whereas the mid-micromolar affinity aligns with a partial shift seen in the analytical SEC. Although this is a relatively low affinity, falls within the range commonly observed for biologically relevant transient interactions^92^.

To confirm that the detected interaction is specific and consistent with the structural model presented above, we produced the ZBTB38^BTB^ D41A mutant to disrupt one of the key predicted contacts between the two proteins. Using the same experimental conditions as for the WT protein, we performed analytical SEC of the mixture of ZBTB38^BTB^ D41A and UBC9, as well as of each protein individually (**Figure 2B, *top***). In this case, the mixture chromatogram displayed better-resolved peaks, of which the first was slightly shifted compared to that for ZBTB38^BTB^ D41A alone, while the second perfectly overlapped with that for UBC9 alone. The accompanying SDS-PAGE analysis showed almost none of the earlier co-elution of the two proteins seen upon mixing the WT proteins (**Figure 2B, *bottom***). This markedly different elution profile demonstrates that the D41A mutation strongly weakened the interaction.

We also produced and analysed the ZBTB38^BTB^ K43R mutant, a conservative substitution replacing the ZBTB38 lysine residue placed inside UBC9’s active site in the AlphaFold3 model with arginine. This mutant was expected to abolish SUMOylation of ZBTB38^BTB^ but have a limited effect on its interaction with UBC9. In line with these expectations, when analysed under the same analytical SEC conditions as above, this mutant did not appreciably affect the ZBTB38^BTB^:UBC9 elution profile compared to that of the WT proteins (**Figure 2C, *top***, albeit with slightly less apparent co-elution on SDS-PAGE, ***bottom***), indicating that the transient interaction is at least largely preserved.

Altogether, these findings support the existence of a direct physical interaction between ZBTB38^BTB^ and UBC9, and are consistent with the binding being mediated by the conserved bottom ZBTB38^BTB^ surface functioning as a structural mimic of the linear SUMO consensus motif. The Asp_41_ residue – which in the AlphaFold3 model serves a role equivalent to that of the +2 glutamate of the consensus motif – is validated as important for this interaction.

### The UBC9-recruiting BTB surface promotes ZBTB38^BTB^ SUMOylation in vitro

We next investigated *in-vitro* SUMOylation of ZBTB38^BTB^, beginning with the wild-type (WT) protein. For this assay, the core SUMOylation machinery components comprising the SAE1:SAE2 hetero-dimer, UBC9, and SUMO1, were recombinantly produced in *E. coli* and purified. In the case of SUMO1, the expressed construct comprised residues 18-97, corresponding to a C-terminally processed form additionally lacking the flexible N-terminal region. SUMOylation reactions were performed at the substrate concentration of 15 µM and monitored over 135 min (**Figure 3A, B**).

**Figure 3.**
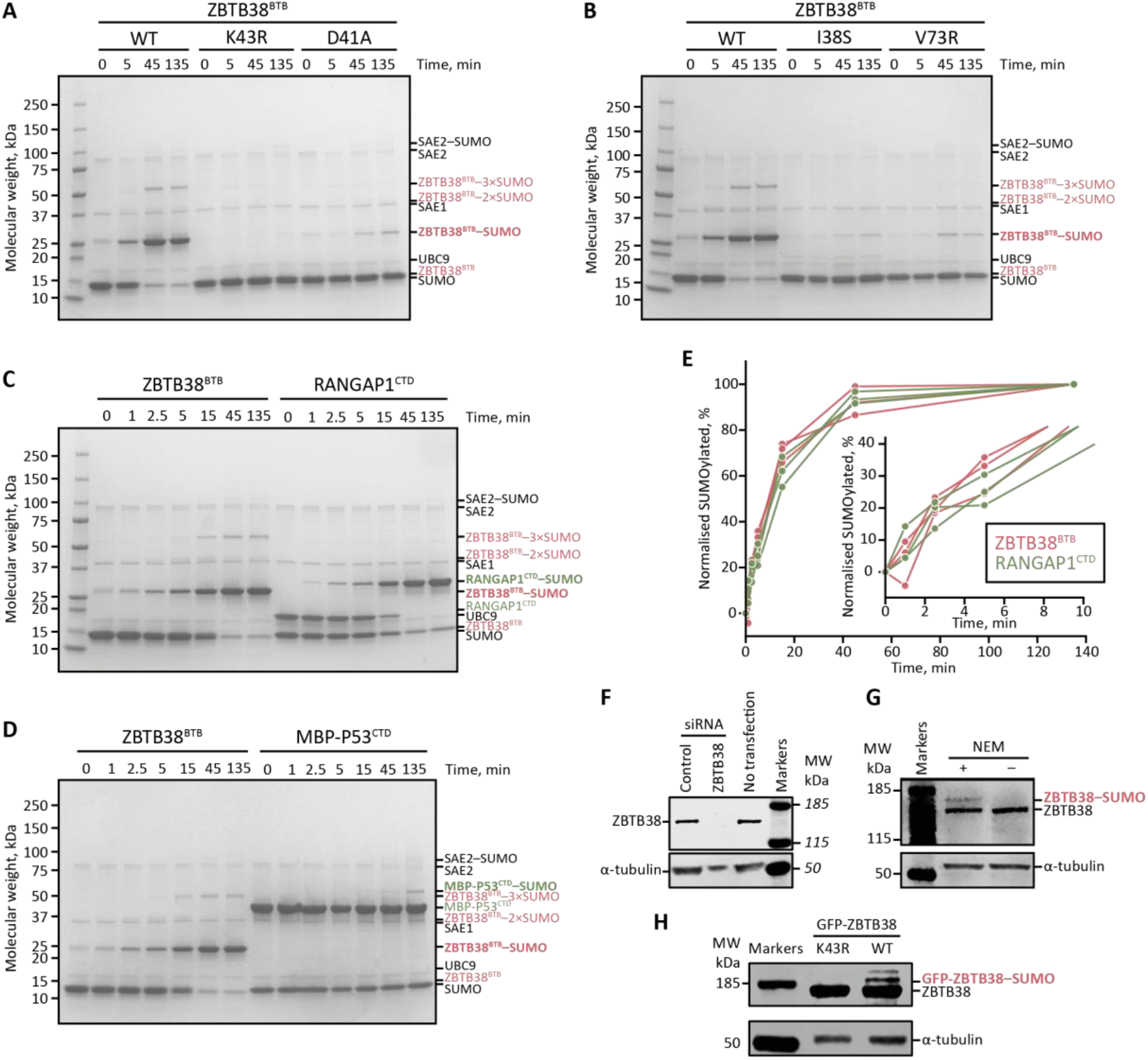
Characterisation of ZBTB38 SUMOylation in vitro and in cells. (A, B) In-vitro SUMOylation assay (15 µM substrate, 0.5 µM E1, 0.5 µM UBC9, 20 µM SUMO1_18-97_, and 5 mM ATP) of indicated ZBTB38^BTB^ variants analysed over time. 0-min time points corresponds to a delay of ∼2 s between ATP addition and reaction quenching. (C, D) In-vitro SUMOylation assays comparing ZBTB38^BTB^ to model substrates, RANGAP1^CTD^ and MBP-P53^CTD^ under the same conditions as in panels A and B. (E) Quantification of mono-SUMOylation-product bands in three technical repeats of a gel-based assay similar to that in panel C, but performed with 20 µM substrate. Inset is a close-up view of the first 10 min of the reaction. (F) Western blot analysis of endogenously expressed full-length ZBTB38 in RPE1 cells transfected with a negative-control or ZBTB38-specific siRNA or untransfected. (G) Western blot analysis of endogenously expressed full-length ZBTB38 in RPE1 cells, with lysates prepared either with or without N-ethylmaleimide (NEM), an inhibitor of cysteine proteases including deSUMOylating enzymes. A probable SUMOylated product is indicated. (H) Western blot analysis of overexpressed green fluorescent protein (GFP)-tagged full-length ZBTB38, WT or K43R, in HEK293T cells rapidly lysed in Laemmli denaturing buffer. Probable SUMOylated products are indicated In F-H, an anti-ZBTB38 (antibody HPA014865) and control anti-α-tubulin western blots are provided.

A band at the apparent MW of 25 kDa corresponding to mono-SUMOylated ZBTB38^BTB^ appeared already at the first time points and intensified over time, culminating in complete SUMOylation of ZBTB38^BTB^ by the end of the experiment (**Figure 3A**; the residual band at 135 min reflects excess SUMO1, which runs at a similar but slightly lower apparent MW, distinguishable upon careful scrutiny of gels). Interestingly, even the 0-min time point, which corresponds to a delay of ∼2 s between adenosine triphosphate (ATP) addition and reaction quenching, reproducibly showed a small amount of mono-SUMOylated product. In addition to the predominant mono-SUMOylation, additional higher bands at the apparent MWs of 37 kDa and 50 kDa, likely corresponding to di-(weak band) and tri-SUMOylated (stronger) ZBTB38^BTB^, respectively, were detected from the 15-min time point onwards. These bands suggest additional, minor SUMOylation events localised to ZBTB38^BTB^ or to the SUMO1 molecule attached to it.

Of the eight lysine residues in our ZBTB38^BTB^ construct, only one, Lys_43_– the residue positioned within the UBC9 active site in our model – has been detected as a SUMOylation site in prior proteomics studies (the other detected ZBTB38 SUMOylation sites fall outside the BTB domain). To experimentally test whether the observed SUMOylation localises to Lys_43_, we carried out SUMOylation of the ZBTB38^BTB^ K43R mutant under the same conditions (**Figure 3A**). Both the predominant mono-SUMOylated and the weaker apparent di- and tri-SUMOylated bands completely disappeared, even at the 135-min time point. Given that the K43R mutation does not affect the ZBTB38^BTB^:UBC9 interaction in the analytical SEC above, the abolished SUMOylation is consistent with Lys_43_ being the major SUMOylation site. Abolishing the apparent multiple SUMOylation events by mutating a single site can be explained by two main scenarios: either i) all ZBTB38^BTB^ WT SUMOylation localises to Lys_43_, with di- and tri-SUMOylated bands representing short SUMO1 chains on that single site; or ii) additional minor SUMOylation sites exist, but they are only modified if Lys_43_ is SUMOylated first. Given that SUMO1 is generally less efficient than SUMO2/3 at forming covalent chains and, additionally, that the main chain-mediating lysine in SUMO1, Lys_7_, is absent from our truncated SUMO1 fragments, we expect no or very little chain formation and thus favour the second scenario, with Lys43-linked SUMOylation promoting further minor ZBTB38^BTB^ SUMOylation events on other lysines, either by releasing SUMO∼UBC9 from its preferred Lys_43_ to access these other sites, or more actively, by recruiting UBC9 through its backside noncovalent SUMO binding site^19^.

To demonstrate that the efficient ZBTB38^BTB^ SUMOylation depends on the modelled interaction between the conserved bottom surface of the BTB domain and UBC9, we used ZBTB38^BTB^ mutants designed to disrupt this interaction based on the AlphaFold3 model: the D41A mutant, shown to impair UBC9 binding in the analytical SEC experiment above, and two single mutants targeting hydrophobic amino acids predicted to interact with hydrophobic pockets of UBC9, Ile_38_ and Val_73_, mutated to a serine and arginine, respectively. With all three mutants, the level of SUMOylation was drastically reduced, with low levels of mono-SUMOylation seen only at longer time points (**Figure 3A, B**). Collectively, these results indicate that the three residues constituting an apparent non-contiguous SUMOylation motif in our model play a critical role in promoting efficient ZBTB38^BTB^ SUMOylation.

### ZBTB38^BTB^ is SUMOylated with comparable efficiency to RANGAP1^CTD^ in vitro

To further evaluate the robust *in-vitro* SUMOylation of ZBTB38^BTB^, we compared it with that of two well-characterised SUMOylation substrates containing a SUMOylation consensus motif: RANGAP1^CTD^, the already introduced uniquely efficient SUMOylation substrate, and a fragment containing the tetramerisation and the C-terminal domains of P53 (referred to here as P53^CTD^), whose linear FKTE motif located in its disordered part^93,94^ undergoes weak SUMOylation on Lys_386_ without a SUMO E3 ligase *in vitro*^95^. P53^CTD^ was tagged at the N-terminus with maltose-binding protein (MBP) to facilitate its expression and purification; the tag was retained in the assay.

We repeated ZBTB38^BTB^ WT SUMOylation under almost the same conditions as above, but over more time points, and compared these results with parallel RANGAP1^CTD^ or MBP-P53^CTD^ SUMOylation kinetics (**Figure 3C, D**). The SUMOylation of RANGAP1^CTD^ was very efficient, both in terms of its initial rate and the fact that the protein became completely modified (**Figure 3C**). Strikingly, ZBTB38^BTB^ is SUMOylated in a similar manner, with the apparent rate throughout the time course being roughly comparable to that of RANGAP1, and with a comparable final completeness (**Figure 3E**). This is remarkable in light of the history of the field, in which no other substrate has been shown to undergo modification comparable to that of RANGAP1^CTD^ under any of the E3-independent *in-vitro* conditions tested. Of note, the MBP-P53^CTD^ case, where the product band only became clearly distinguishable at the 135-min time point (**Figure 3D**), provides a dynamic-range control for our experiment, illustrating that under the conditions used, the SUMOylation reaction can be strongly limited for certain substrates, and making the highly efficient modification of ZBTB38^BTB^ all the more noteworthy.

### Potential cellular SUMOylation of ZBTB38

To investigate ZBTB38 SUMOylation in a cellular context, we first sought to identify a human cell line expressing the endogenous protein at detectable levels. Mining public expression databases showed that ZBTB38 mRNA was abundant in RPE1 cells, a human retinal pigment epithelial cell line that is frequently used as a model of non-transformed cells. Western blotting revealed the presence of a protein at the expected MW cross-reacting with an anti-ZBTB38 antibody (∼134 kDa, **Figure 3F**). Transfection of siRNA targeting ZBTB38 or a negative control ascertained the specificity of the band detected. Therefore, we conclude that RPE1 cells express endogenous ZBTB38 at detectable levels.

Because SUMO modification is sensitive to SUMO-specific SENP proteases present in cells, we performed extraction both by standard procedure and in the presence of N-ethylmaleimide (NEM), a thiol-alkylating agent that inhibits cysteine proteases, including SENPs (**Figure 3G**). While no higher-MW band was detectable above the main ZBTB38 band with an anti-ZBTB38 antibody under standard extraction conditions, at least one such higher band was observed with NEM-supplemented extraction. Based on the apparent MW and NEM-dependence, this band likely corresponds to a ubiquitin-like modification, such as SUMO.

To shed light on the localisation of this modification, we turned to human embryonic kidney (HEK) 293T cells, in which endogenous ZBTB38 is undetectable, thereby facilitating experiments performed with exogenous proteins. Cells were transfected with plasmids encoding full-length green fluorescent protein (GFP)-tagged human ZBTB38 in either a WT or a K43R mutant variant. In this case, to avoid deSUMOylation by SENPs, we resorted to rapid lysis under denaturing conditions rather than NEM supplementation. Under these conditions, cells overexpressing WT ZBTB38 showed a set of higher-MW bands with the anti-ZBTB38 antibody, including, again, one stronger species (**Figure 3H**). In contrast, the K43R mutation abolished all these additional bands, indicating that the observed modification is dependent on, and probably localised to, Lys_43_. The apparent dependence of the identified modification specifically on Lys_43_– the residue positioned inside the UBC9 active site in our structural model and required for ZBTB38^BTB^ SUMOylation *in vitro* – argues that this modification is SUMOylation.

Taken together, these observations are consistent with ZBTB38 SUMOylation, thus extending our *in-vitro* results to a cellular context using a full-length, and partly endogenous, protein. We note the congruence of these observations with those previously obtained for both endogenous and overexpressed forms of a close homologue of ZBTB38, ZBTB33, by Zhenilo, Prokhortchouk and colleagues, who went on to confirm that the ZBTB33 modification corresponds primarily to SUMO1 and is dependent on Lys_42_ (equivalent to Lys_43_ of ZBTB38)^59^.

### BTB domains of five ZBTB proteins share UBC9-interaction site and SUMOylation propensity

To complete our study, we investigated whether the interaction with UBC9 and the resultant efficient SUMOylation are more frequent features of BTB domains of ZBTB proteins beyond ZBTB38. We already noted that the identified UBC9-interacting surface is conserved within one clade of ZBTB proteins including ZBTB4, ZBTB14, ZBTB21, ZBTB33, and ZBTB38, and that all of these proteins have been reported to be SUMOylated on the expected lysine residue in human cells^28,33,59,61,77^. Here we wanted to revisit this question in an unbiased manner.

To identify potential binding partners of UBC9 among all human ZBTB^BTB^ domains *in silico*, we performed a computational protein:protein interaction screen using our custom AlphaFold3-based framework introduced above. This time, UBC9 was used as a bait, while the query set comprised the 51 BTB domains of all human ZBTB-family proteins and two ZBTB-like proteins, NACC1 and NACC2. Each BTB domain was repeated twice in tandem to obtain a single-chain, fused BTB dimer-like arrangement, like done above for the ZBTB38^BTB^ interactor screen. This screening recovered the entire phylogenetic clade of ZBTB38 among the top-ranked predicted UBC9 interactors, with ZBTB21^BTB^ and ZBTB33^BTB^ as the highest-scoring candidates (**Figure 4A**). The only ZBTB protein from outside this clade whose BTB domain scored comparably was ZBTB47^BTB^, but visual inspection of obtained models indicated that it is predicted to interact *via* a different surface and without inserting a lysine residue into the active site of UBC9. Whether this prediction represents a real interaction needs to be investigated in the future, but structurally the predicted ZBTB47^BTB^:UBC9 interaction does not belong to the same class as the others.

**Figure 4.**
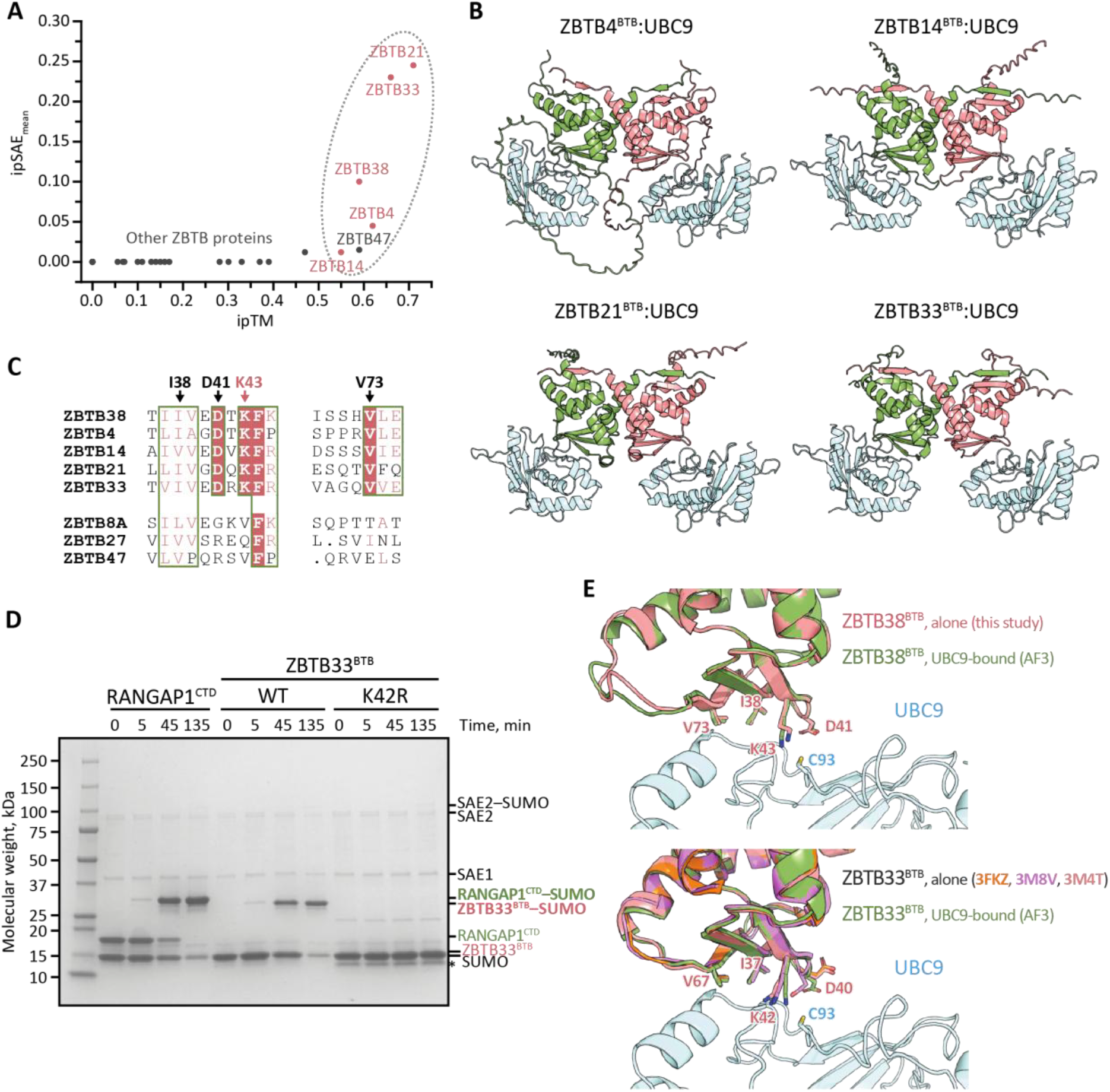
Conservation of UBC9-interaction and SUMOylation propensities in other ZBTB proteins. (A) Scatter plot representing the results of computational screening of candidate ZBTB^BTB^ dimers against UBC9, plotted according to ipTM and ipSAE_mean_ metrics. (B) AlphaFold3 models of the ZBTB^BTB^:UBC9 complexes in 2:2 stoichiometry. (C) Fragments of a multiple sequence alignment of selected ZBTB proteins from the ZBTB38 clade (top) and outside it (bottom). The sequence fragments correspond to the UBC9-interacting surface, with the target Lys_43_ and the key interface residues labelled using ZBTB38 numbering. (D) In-vitro SUMOylation assay comparing ZBTB33^BTB^ WT with its K42R mutant and with RANGAP1^CTD^ under the same conditions as in 3A and B. The thin UBC9 band (around 17 kDa) is left unlabelled to avoid crowding. The lowest band in K42R wells, marked with an asterisk, corresponds to a contaminant. (E) Structural alignment of crystal structures of ZBTB38^BTB^ (top) and ZBTB33^BTB^ (bottom) with their respective UBC9-bound AlphaFold3 (AF3) models, all in cartoon representation. Selected interface residues on the BTB domains and the catalytic Cys_93_ on UBC9 are shown as sticks. For ZBTB33^BTB^, the indicated codes are Protein Data Band (PDB) accessions.

For the ZBTB38-clade proteins, all BTB domain dimers were modelled to interact with UBC9 in essentially the same manner through their conserved bottom surface, with the equivalent SUMO acceptor lysine (Lys_40_ of ZBTB4, ZBTB21; Lys_42_ of ZBTB33; Lys_43_ of ZBTB38; Lys_46_ of ZBTB14), well-positioned within the UBC9 active site, close to the catalytic cysteine. This arrangement was preserved when each domain was remodelled as a dimer bound to UBC9 with a corrected, 2:2, stoichiometry (**Figure 4B**). In line with the ConSurf mapping presented above, sequence alignment of the two non-contiguous ZBTB38 regions involved in UBC9 interaction (residues 37–45 and 70–75) with the corresponding regions of selected other ZBTB proteins revealed that all key predicted interface residues are conserved across the other ZBTB38-clade members but generally not in other ZBTB proteins (**Figure 4C**). Together, these observations indicate that the UBC9-binding interface is a characteristic feature of the ZBTB38-clade proteins that was likely already present in their common ancestor and has seemingly been under strong evolutionary constraint across the whole clade.

Based on these structural observations, we next investigated the SUMOylation of ZBTB33^BTB^, to evaluate whether it is similarly efficient to that of ZBTB38^BTB^. SUMOylation of ZBTB33 has been studied before in cells^59^, but has never been reconstituted *in vitro*. In addition to the WT domain, we generated a mutant in which the expected target Lys_42_ residue was substituted with arginine (ZBTB33^BTB^ K42R). ZBTB33^BTB^ was SUMOylated with E1 and E2 enzymes under equivalent conditions to those used for ZBTB38^BTB^ above, and compared with RANGAP1^CTD^(**Figure 4D**). In agreement with the results obtained for ZBTB38^BTB^, a band with an apparent MW of 25 kDa, corresponding to mono-SUMOylated ZBTB33^BTB^, was readily detectable within minutes. Under the conditions used, the WT ZBTB33^BTB^ undergoes SUMOylation apparently as efficiently as RANGAP1^CTD^, and thus probably as WT ZBTB38^BTB^. However, unlike for ZBTB38^BTB^, no higher-MW bands corresponding to di- or tri-SUMOylation were observed for this substrate. In contrast to the WT, the ZBTB33^BTB^ K42R mutant showed no SUMOylation at all, indicating that Lys_42_ residue is the only SUMOylation site within this domain. In the previous experiments performed in cells, full-length ZBTB33 showed near-complete SUMOylation upon overexpression of SUMO1, SUMO2, or SUMO3^59^, a striking observation given the usual low SUMOylation levels detected on substrates in cells, even when overexpressing SUMO; these previous results are consistent with the exceptional intrinsic SUMOylability of ZBTB33^BTB^ and ZBTB38^BTB^ demonstrated here.

Overall, structural predictions, comparative structural and sequence analyses, and *in vitro* SUMOylation assays of two clade members, ZBTB38 and ZBTB33, provide strong evidence that the entire set of BTB domains within this phylogenetic clade – including ZBTB4, ZBTB14, ZBTB21, ZBTB33, and ZBTB38 – interact with UBC9 through the same conserved surface and, in consequence, undergo efficient SUMOylation on a conserved lysine residue.

### The BTB SUMOylation motif is pre-organised for UBC9 binding and SUMOylation

To provide a tentative explanation for the unusually efficient SUMOylation of this set of BTB domains, we reasoned that their UBC9-binding surfaces and target lysine residues might be spatially pre-organised in a suitable conformation for binding and reaction, thereby accelerating the substrate recruitment and lysine positioning steps of the SUMOylation cascade, which are normally rate-limiting, particularly for a conformationally flexible linear SUMOylation motif such as that in P53^CTD93,94^. Such a mechanism would minimise the entropy penalty of UBC9 binding and productive lysine positioning, and, in statistical terms, increase the probability both of a productive encounter between a BTB domain and UBC9 and correct alignment of the reacting lysine and cysteine residues.

To test this hypothesis, we scrutinised the spatial arrangement of all interface residues by super-imposing the predicted ZBTB38:UBC9 complex onto our experimentally determined structure of ZBTB38^BTB^. In parallel, we similarly aligned the predicted ZBTB33:UBC9 complex with the three previously determined ZBTB33^BTB^ structures deposited in the PDB by other teams (3M4T, 3M8V, and 3FKC, all lacking accompanying publications). All analysed experimental BTB domain structures super-impose closely onto the corresponding domain in the predicted UBC9-bound complexes (**Figure 4E**). Indeed, in addition to the overall three-dimensional superposition of secondary-structural elements, the side-chains of individual residues – particularly the SUMOylated lysine and others directly involved in the interaction with UBC9 – are closely aligned between the experimental UBC9-free and the predicted UBC9-bound models, with a partial exception of Asp_40_ in ZBTB33 (equivalent to Asp_41_ in ZBTB38), which alternates between two distinct rotamers across *apo* ZBTB33^BTB^ structures. Furthermore, we note that prior nuclear magnetic resonance (NMR) data have shown that the BTB domain of ZBTB33 – unlike that of ZBTB17/MIZ1 – is conformationally rigid^96^ and would thus be expected to undergo little conformational change upon partner binding, particularly when that binding is of relatively low affinity. These observations are consistent with our pre-organisation hypothesis.

## Discussion

In this study, we determined the first three-dimensional X-ray crystallography structure of ZBTB38^BTB^ and identified the structural basis underlying its modification with an essential eukaryotic protein PTM, SUMOylation. Our analyses suggest the existence, within this protein, of an atypical SUMOylation motif that directly recruits the SUMO-conjugating E2 enzyme UBC9, enabling efficient modification of the central lysine, Lys_43_, in the absence of a SUMO E3 ligase. SUMOylation of full-length ZBTB38 on the same site has been previously evidenced in cells using high-throughput proteomics^28,77,33,60^, and we show, for the first time, higher-MW bands consistent with appreciable SUMOylation for both endogenous and overexpressed ZBTB38 under endogenous levels of SUMO cascade components in human cells. We further show that UBC9-recruitment and efficient SUMOylation are not unique to ZBTB38^BTB^, but are likely shared by BTB domains of a vertebrate ZBTB-protein clade additionally including ZBTB4, ZBTB14, ZBTB21, and ZBTB33 across species. This provides the biochemical basis for previous reports of functionally important ZBTB33 and ZBTB14 SUMOylation on an equivalent lysine in human cells and fish, respectively^59,61^, in addition to informing potential future studies on the functional impact of SUMOylation of ZBTB38 and remaining clade members.

Our characterisation of the ZBTB38^BTB^ oligomeric state in solution by SEC (**Figure 2A-C**) and our determination of its three-dimensional structure by X-ray crystallography (**Figure 1B**) demonstrate that this domain is predominantly dimeric. This is unsurprising given the available data on other BTB domains, but it bears on past controversies about the contribution of the BTB domain to ZBTB38 self-association in cells^97^. Homo-dimerisation is once again confirmed as a fundamental structural feature of canonical ZBTB-type BTB domains and their primary, conserved function (**Figure 5A**).

**Figure 5.**
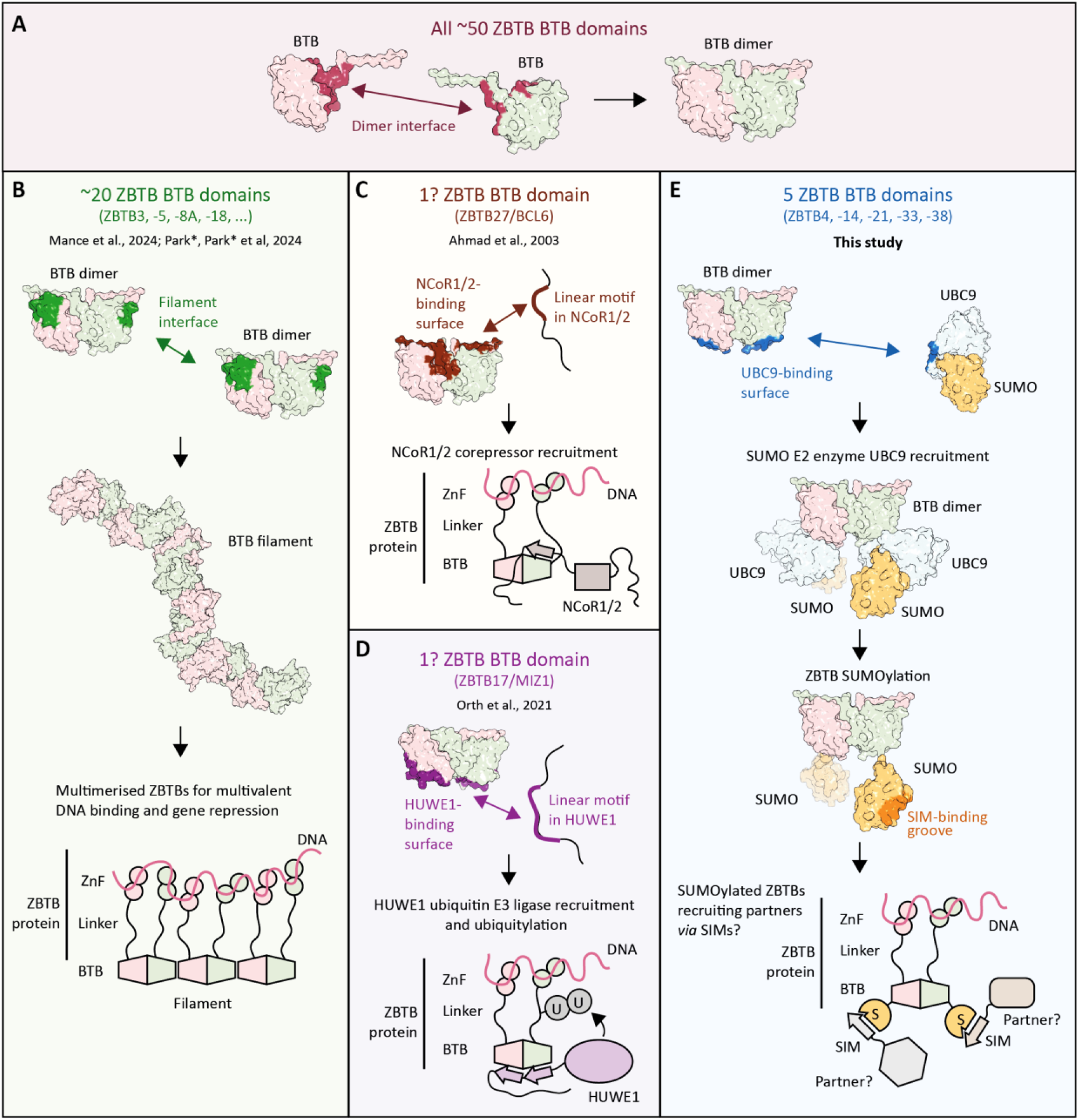
Molecular functions of BTB domains of ZBTB transcription factors. (A) The BTB domains of all ∼50 human ZBTB proteins likely homo-dimerise through a conserved dimer interface. (B) The BTB domains of ∼20 ZBTB proteins further self-assemble via a filament-forming inter-dimer interface to support multivalent DNA binding and gene repression. (C) The BTB domain of ZBTB27/BCL6 presents a surface that binds a linear motif in the NCoR1/2 co-repressors. (D) The BTB domain of ZBTB17/MIZ1 has a distinct surface that binds a linear motif present in the HUWE1 ubiquitin E3 ligase, promoting ubiquitylation. (D) Our study identifies the BTB domains of five human ZBTB proteins as UBC9-recruiting platforms, leading to efficient ZBTB SUMOylation. The resulting SUMO marks may in turn recruit downstream partners through their SUMO-interacting motifs (SIMs).

A further function, reported or predicted for a total of twenty or 40% of human ZBTB-family members, involves higher-order assembly of dimers into filaments^71,73^ (**Figure 5B**). In our previous study on filament-forming ZBTB BTB domains, we reported filament formation to depend on a specific interdimer interface^73^. While residues involved in this interface are partly different for each filament-forming ZBTB protein – possibly to favour homo-multimerisation of each independently of others – almost all filament-forming proteins appear to share a conserved serine residue around position 112 (using ZBTB38 numbering), which is located at the narrow bottleneck of the filament interface. Mutation of this residue to a bulkier side chain has been shown to abolish filament formation in all tested cases. In ZBTB38^BTB^, this residue is Lys_112_. The presence of this bulky and positively charged residue is therefore incompatible with the formation of filaments similar to those observed by us and others for selected ZBTB^BTB^ domains. Although other, independently evolved higher-order interfaces are conceivable, and some have been postulated based on crystal contacts (without conclusive evidence in solution) for several ZBTB^BTB^ domains^79–82^, our experimental results (SEC, crystal contact analysis, AlphaFold3 modelling of multimers) do not support any strong tendency of ZBTB38^BTB^ to form assemblies beyond the canonical dimer in solution (**Figure 2A-C, Supplementary Figure 1**).

Instead, our identification of a conserved, UBC9-interacting surface on ZBTB38^BTB^ reveals that, rather than mediating filamentation, this and related BTB domains might serve as particularly efficient SUMOylation hubs. Thus, alongside the already mentioned homo-dimerisation (**Figure 5A**) and filamentation (**Figure 5B**) propensities, and previously identified NCoR1/2-recruitment^79^ (**Figure 5C**) and HUWE1-recruitment^98^ (**Figure 5D**) functions, UBC9 recruitment through the identified surface emerges as the new molecular function of BTB domains (**Figure 5E**).

The revealed BTB-domain SUMOylation site is distinguished from typical SUMO sites by its structured character, and the way a spatial arrangement of residues mimics a canonical linear SUMOylation motif (ΨKXE or its variants) to engage UBC9 (**Figure 1G**). Superficially, Lys_43_ lies within one of the variant SUMOylation motifs: the inverted SUMOylation motif E/DXKΨ. However, structural modelling of the ZBTB38^BTB^:UBC9 complex indicates that the expected hydrophobic residue located in the +1 position relative to the SUMOylated lysine, Phe_44_, is oriented away from the solvent and towards the hydrophobic core, thus not contributing to the UBC9 interface. Instead, van der Waals interactions are mediated by sequentially more distant residues, including Val_73_ and Ile_38_, which, together with Asp_41_, form a structural SUMOylation motif surrounding Lys_43_ (**Figure F**). This interpretation is supported by the drastic decrease in SUMOylation efficiency (**Figure 3A-B**) and in interaction with UBC9 (**Figure 1A-B**) upon individual mutagenesis of these residues. Moreover, the residues in question are conserved, alongside the modified lysine, across the ZBTB38 clade (**Figure 4C**). Given the possibility of structural SUMOylation motifs not readily detectable from sequence revealed in this study, and previously for UBC9 itself^27^, UBE2K^23^, and NF1^53^, the approach used by us here – namely computational detection of UBC9 binders *via* a high-throughput AlphaFold3 screen – might represent an interesting complementary approach *vis-à-vis* sequence-based SUMOylation motif detection, with the latter valid for disordered linker or loop regions of substrates but generally not for folded domains.

The affinity of ZBTB38^BTB^ for UBC9 measured with ITC in this study (∼65 µM, **Figure 2D**) is in the same mid-micromolar range as the affinities determined with the same technique for peptides derived from four established SUMOylation substrates: RANGAP1 (∼13 µM), ELK1 (∼70 µM), CBP (∼55 µM), and Calpain 2 (∼55 µM)^99^. The *K*_D_ of the complete RANGAP1^CTD^ domain has not, to our knowledge, been measured, but it may be comparable, judging by its SUMOylation *K*_M_ value in the mid-micromolar range^26^. By contrast, peptides derived from P53 and JUN have been shown, albeit using a different technique (NMR), to bind UBC9 far more weakly, with *K*_D_ in the low-millimolar range, and we expect that many other substrates bind UBC9 similarly poorly, explaining the scarcity of *K*_D_ measurements in the literature. The affinity of ZBTB38^BTB^ is therefore probably above average, and this may partly explain why its SUMOylation is efficient. Of note, micromolar affinities might appear weak in absolute terms, but similar values are commonly observed for PTM and other enzyme:substrate systems, where they may keep activity tunable by mechanisms that transiently raise the effective local substrate concentration, such as co-localisation, scaffolding, or multivalency^92,100,101^.

The affinity of ZBTB38^BTB^ for UBC9, while possibly above average, is nonetheless not outstanding, which may have the functional advantages mentioned above but means that the exceptional efficiency and completeness of its SUMOylation seen in our *in-vitro* assays require a further explanation. SUMOylation is a complex, multi-step process, and substrate recruitment to UBC9 is only one of its steps. While the modification of many substrates, especially at low concentrations, may be limited largely by inefficient substrate recruitment, RANGAP1^CTD^ is known to differ from other substrates not only by a lower *K*_M_– as would be expected if substrate recruitment was the sole differentiating factor – but also by a higher *k*_cat_ and a higher final modification yield (complete rather than partial modification)^102^. We suspect that a key determinant of efficient SUMOylation – possibly affecting *k*_cat_ in particular – is the positioning of the target lysine at the active site of the loaded UBC9, which in turn – as discussed below – might be related to the structural pre-organisation of the substrate. In our assays, we used substrate concentrations (15-20 µM) that should be sub-saturating for ZBTB38^BTB^ and possibly also for RANGAP1^CTD^, and an elevated E1 concentration (0.5 µM) that ensures efficient UBC9 recharging, allowing us to focus on the final, UBC9-dependent SUMOylation steps. Under these conditions, ZBTB38^BTB^ showed the apparent modification rate similar to that of RANGAP1^CTD^ throughout the time course, pointing to comparable apparent catalytic efficiency (*k*_cat_/*K*_M_) for both substrates (**Figure 3C,E**). Furthermore, the high completeness of ZBTB38^BTB^ SUMOylation indicates that its modification rate did not collapse as free substrate was gradually depleted and product accumulated, as for RANGAP1^CTD^, but unlike what is typically observed for other substrates, even otherwise relatively efficient ones such as TDG^25^. We note that our analysis is qualitative, and the exact ZBTB38^BTB^ SUMOylation kinetics, and how comparable they are to those for RANGAP1^CTD^, are likely sensitive to experimental conditions, including substrate and enzyme concentrations and the presence or absence of an E3 ligase (absent here). Nonetheless, although assay conditions vary across the field, this is, to our knowledge, the first time that SUMOylation comparable to that of RANGAP1^CTD^ in efficiency and completeness has been observed for any other substrate under E3-less conditions *in vitro*, indicating an exceptional intrinsic SUMOylability of ZBTB38 likely relevant for its biological function.

Proteomic and biochemical studies have shown that SUMO acceptor lysines are often located within flexible regions, such as intrinsically disordered regions or exposed loops^33,34,103^. Such locations likely facilitate UBC9’s access to target lysines.

However, flexibility alone does not guarantee efficient SUMOylation and, indeed, could be detrimental to it. Excessive local dynamics may reduce the probability of adopting a productive binding and catalytic geometry, increasing the entropic cost of productive substrate engagement and resulting in inefficient SUMOylation (**Figure 6, *left***). In this context, we suggest BTB domains are exceptional SUMO substrates in part precisely because their UBC9-binding surface and acceptor lysine are locked into conformations that are close to those adopted in the bound and reaction-proficient state (**Figure 6, *right***). As the mid-micromolar affinity of ZBTB38^BTB^ for UBC9 apparently does not exceed that possible for stronger-binding linear peptides, we suspect that the main advantage of the structural BTB site lies in its correct pre-organisation for the SUMOylation reaction itself rather than for UBC9 binding, although the latter might play a role as well. Unlike a flexible linear motif such as P53^CTD^’s, which must be re-organised at an entropic cost, a rigid domain pays that cost up front, thereby reducing the conformational penalty associated with recruitment and lysine positioning. Consistent with this model, experimental UBC9-free structures of ZBTB38^BTB^ and ZBTB33^BTB^ super-impose closely on their predicted UBC9-bound forms, including much of the acceptor lysine side-chain (**Figure 4E**), indicating little adaptation is required to engage the enzyme and adopt a position conducive towards the nucleophilic attack required for SUMO transfer. The previous NMR evidence that the ZBTB33^BTB^ domain is highly intrinsically rigid^96^ provides additional support in favour of the pre-organisation model. A similar mechanism might in part explain the high SUMOylation efficiency of RANGAP1^CTD^, as its consensus motif is located on a constrained loop protruding from a folded domain rather than a disordered region.

**Figure 6.**
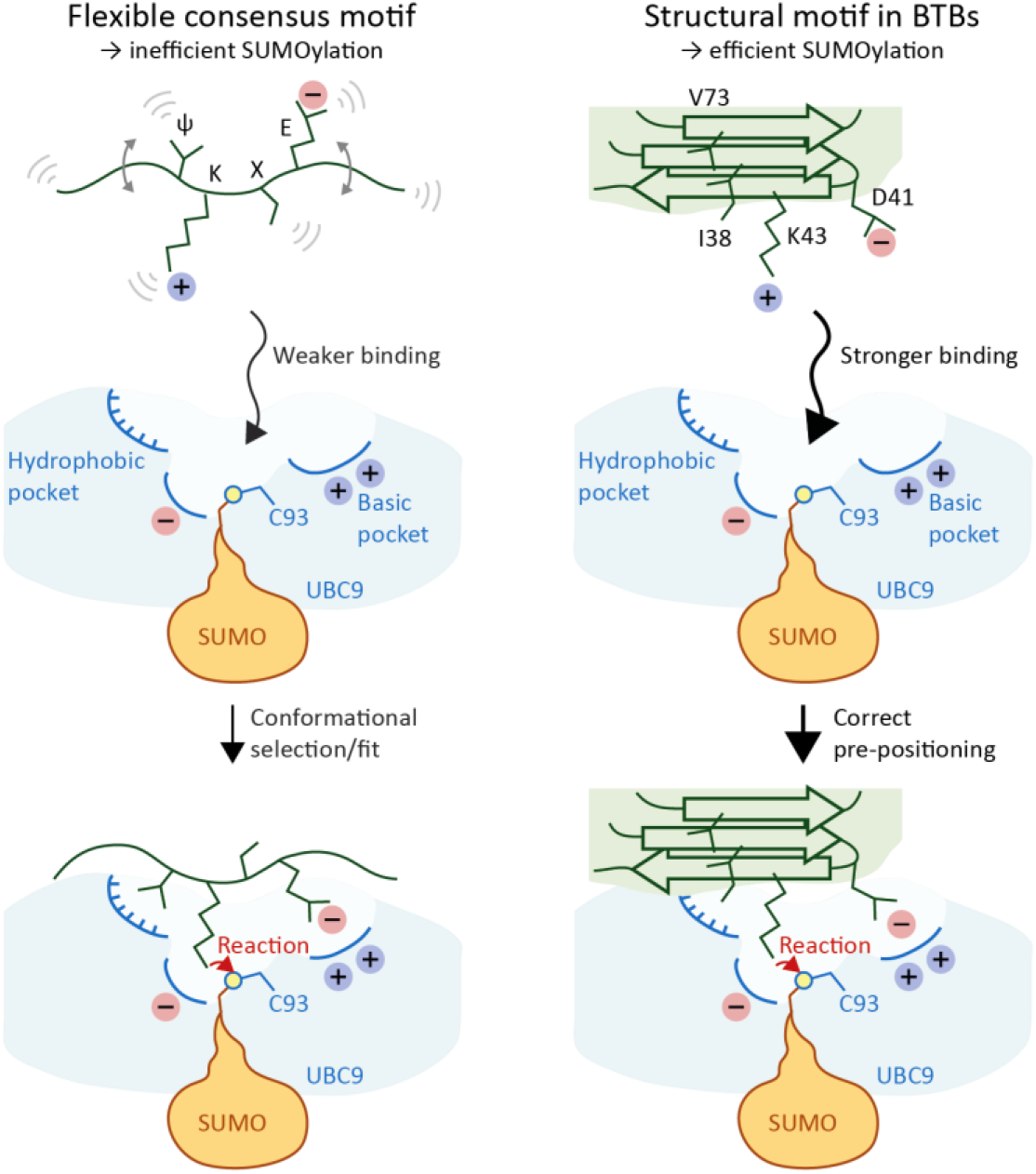
Proposed mechanism for efficient BTB-domain SUMOylation. Canonical SUMOylation relies on a flexible linear consensus motif (ΨKXE) that binds UBC9 weakly and must be selected or induced into a reactive conformation, making SUMO transfer from the catalytic cysteine Cys_93_ onto the lysine inefficient. By contrast, in ZBTB^BTB^ domains, the equivalent lysine (e.g. Lys_43_ of ZBTB38) and spatially proximal UBC9-binding residues are presented on a rigid, folded structural element. This pre-organised motif binds UBC9 more strongly and is correctly pre-positioned relative to Cys_93_, enabling efficient SUMO transfer.

While SUMOylation can be enhanced in cells by SUMO E3 ligases, we expect that intrinsic SUMOylability at the level of the E1 and E2 enzymes – such as that revealed here for ZBTB^BTB^ domains – generally serves as the basis for such enhancement, thus playing a decisive role in the relative modification levels of different sites even where E3 ligases are present. Indeed, to date only a single substrate for which an E3 ligase overrides the intrinsic site specificity of UBC9 has been documented (PCNA^102,104,105^); in all other cases studied so far, E3 ligases appear to enhance UBC9’s activity, and possibly its co-localisation with substrates, but not change its site preference. High intrinsic SUMOylability might be particularly critical for substrates that are robustly SUMO1-modified under basal, non-stressed conditions, such as RANGAP1 or, indeed, ZBTB33^59,106^. Thus, although SUMO2/3-dependent SUMOylation of ZBTB4, ZBTB14, ZBTB21, and ZBTB38 on residues equivalent to ZBTB38’s Lys_43_ has been detected in cells with mass spectrometry – for which SUMO2/3-specific protocols have been developed more extensively than SUMO1-specific ones – we hypothesise that endogenous forms of these proteins might be primarily SUMO1-modified, by analogy with ZBTB33, for which SUMO1 appears to be the preferred modifier under endogenous conditions^59^, and in line with the notion that SUMO1 is the dominant SUMO paralogue under non-stressed conditions^106^. Indeed, at least one of the sites discussed here, Lys_40_ of ZBTB21, has been detected in one of the small available SUMO1 proteomics datasets^29^. The predominant SUMO1 modification anticipated in cells could also partly explain why the SUMO2/3 intensities on ZBTB peptides detected with mass spectrometry in cells, while relatively high (ZBTB38 Lys_43_ and ZBTB4 Lys_40_ are close to the top 400 or 1% of all detected sites in intensity in a large semi-quantitative study^33^), are not as high as perhaps could be expected based on their *in-vitro* efficiency. Intriguingly, although best supported, by a dedicated cellular study^59^, ZBTB33 Lys_42_ is the only SUMOylation site in this clade not detected by SUMO2/3 proteomics in cells, most probably because it happens to lie within an extremely short tryptic fragment, RK_mod_FR (where ‘mod’ is the expected SUMO modification remnant). SENPs are another key factor shaping the steady-state SUMOylation levels and, to some extent, the SUMO1 *vs* SUMO2/3 specificity, in cells; this aspect of ZBTB SUMOylation regulation remains to be separately investigated.

Lastly, while our study does not address the biological relevance of the uncovered UBC9 interaction and SUMOylation propensity of ZBTB38, we note that the residues involved in this process are under apparent strong evolutionary constraint in the whole ZBTB38 clade (**Figures 1C** and **4C**). This indicates important function(s), whose details may vary across clade members and specific contexts, but which, at a molecular level, likely involve recruitment of SUMO-interacting motif (SIM)^107^-containing factors to SUMOylated ZBTBs (**Figure 5E**). For ZBTB33 and ZBTB14, functional relevance has already been demonstrated, albeit not in precise molecular detail. In the case of ZBTB14 in the zebrafish model, Lys_40_SUMOylation was shown to be essential for ZBTB14-mediated repression of the PU.1 transcription factor, which in turn is necessary for the correct development of monocytes and macrophages^61^. In the case of ZBTB33, abolishing BTB SUMOylation through K42R mutation led to global changes in transcription, with ∼500 downregulated and ∼500 upregulated genes in a human cell line^59^. This suggests a complex picture where ZBTB33 SUMOylation might impact – whether directly or indirectly – both activation and repression of genes. Of note, recent large-scale scanning of human transcription factor sequences by the Bintu laboratory found that linear SUMOylation consensus motifs are by far the most strongly enriched sequence features in regions with repressive tendency^108^ (note that this analysis would miss structured SUMOylation sites such as those described here in ZBTB proteins), suggesting a high potential for SUMOylation in gene regulation. In that light, a previous observation that the mouse ZBTB38^BTB^ domain is sufficient for repression of a luciferase reporter might be linked to its high SUMOylability^109^. Beyond simple transcriptional function, ZBTB38 and two members of its clade, ZBTB4 and ZBTB33, are the only ZBTB proteins so far shown to specifically recognise methylated DNA^76,110,111^. As all three proteins contain the conserved BTB SUMOylation site described here, it is possible that SUMOylation is linked to their roles as readers and downstream effectors of DNA methylation marks.

These different functional possibilities remain to be explored in future studies. Furthermore, the determinants of inherently efficient SUMOylation revealed here may be applicable more broadly across the SUMOylation field, and could facilitate the search for other efficient substrates and the design of SUMOylation decoys and modulators.

## Limitations

Our study has several limitations that suggest avenues for further work. First, although the proposed UBC9-bound model is strongly supported by several lines of evidence (the moderate confidence scores, the robustness of AlphaFold3 modelling across seeds and across ZBTB-clade members, structurally convincing features such as the positioning of the target lysine within the UBC9 active site, and the consistent results of model-informed mutagenesis), it would benefit from validation by an experimental structure. Our extensive attempts to co-crystallise ZBTB38^BTB^ with UBC9 have been unsuccessful, but testing BTB domains from other clade members may allow successful co-crystallisation. Second, while the strict dependence of the observed SUMOylation on Lys_43_ of ZBTB38, and on the equivalent Lys_42_ of ZBTB33, is best explained by these residues being the sites of modification – particularly as the K43R mutant of ZBTB38 largely retains UBC9 binding – we cannot fully exclude an indirect effect, and the modification site could be mapped by complementary methods to ensure proper localisation. We note, however, that SUMOylation of Lys_43_ of ZBTB38, and of the equivalent lysines in most other clade members, has been reproducibly detected in cells by mass spectrometry. Third, our gel-based kinetic experiments should be interpreted as a relative measure of SUMOylation efficiency, constrained by the use of single reaction conditions, including fixed enzyme and substrate concentrations. Although the modification efficiency is closely comparable for RANGAP1^CTD^ and ZBTB38^BTB^ throughout our time course, the two substrates may arrive at this similarity through different mechanisms, e.g. different combinations of the *K*_M_ and *k*_cat_ components. It is also conceivable that the two proteins share a common kinetic ceiling that ‘hides’ further differences between them, although our use of elevated E1 concentration (0.5 µM, 1:1 with UBC9) should ensure efficient recharging of UBC9, moving the kinetic bottleneck downstream, to one of the UBC9- and substrate-dependent processes. Regardless of these caveats, this result places both ZBTB38^BTB^ and RANGAP1^CTD^ in a highly efficient class that is clearly distinct from, for instance, P53^CTD^. Finally, our cellular assays are consistent with appreciable Lys_43_-localised SUMOylation of ZBTB38 but do not unambiguously demonstrate it, as the identity of the bands detected in cells remains to be directly confirmed.

## Materials and Methods

### Plasmids and mutagenesis

The plasmids encoding SUMO1_18-97_ protein (residues 18-97 of the UniProt sequence P63165|SUMO1_HUMAN), human UBC9/UBE2I (UniProt P63279|UBC9_HUMAN), the full-length human SAE1:SAE2 hetero-dimer (UniProt sequences Q9UBE0|SAE1_HUMAN and Q9UBT2|SAE2_HUMAN) and C-terminal domain of RANGAP1 (RANGAP1 CTD, residues 419-587) were generated by us previously^90,112^ and are available at Addgene (plasmid numbers 228951, 228949, 228950, and 228947).

The codon-optimised (for *Escherichia coli*) DNA sequence encoding the BTB domain of human ZBTB38 (residues 1-134 of the Uniprot sequence Q8NAP3| ZBT38_HUMAN) was synthesised by GeneCust and cloned into the pET-24a, conferring a non-cleavable C-terminal LEHHHHHH tag. Then this vector was used as template to produce four distinct ZBTB38 mutants (I38S, D41A, K43R, and V73R) using a standard mutagenesis PCR, followed by DpnI treatment. The protein sequence with the tag is shown below:

MTVMSLSRDLKDDFHSDTVLSILNEQRIRGILCDVTIIVEDTKFKAHSNVLAASSLYFKNIFWSHTICISSHVLELDDLKAEVFTEILNYIYSSTVVVKRQETVTDLAAAGKKLGISFLEDLTDRNFSNSPGPYLEHHHH HH

The codon-optimised (for *E. coli*) DNA sequences coding for BTB domain of human ZBTB33 in WT and K42R variants (residues 1-134 of the Uniprot sequence Q86T24|KAISO_HUMAN) were synthesised by GeneCust and cloned into the pET-28a vector conferring a non-cleavable C-terminal LEHHHHHH tag.

The codon-optimised (for *E. coli*) DNA sequence encoding a C-terminal P53 fragment (domains TD and CTD, residues 322-393 of the UniProt sequence: P04637|P53_HUMAN) fused to an N-terminal His-tag and MBP tag (residues 29-392 of P0AEX9|MALE_ECOLI) was synthesised by GeneCust and cloned into a pET-28a vector. Although a thrombin cleavage site was present between MBP and P53, it was not used during purification. The final protein sequence with the tag is shown below, with the P53 part underlined.

MHHHHHHEEGKLVIWINGDKGYNGLAEVGKKFEKDTGIKVTVEHPDKLEEKFPQVAATGDGPDIIFWAHDRFGGYAQSGLLAEITPDKAFQDKLYPFTWDAVRYNGKLIAYPIAVEALSLIYNKDLLPNPPKTWEEIPALDKELKAKGKSALMFNLQEPYFTWPLIAADGGYAFKYENGKYDIKDVGVDNAGAKAGLTFLVDLIKNKHMNADTDYSIAEAAFNKGETAMTINGPWAWSNIDTSKVNYGVTVLPTFKGQPSKPFVGVLSAGINAASPNKELAKEFLENYLLTDEGLEAVNKDKPLGAVALKSYEEELVKDPRIAATMENAQKGEIMPNIPQMSAFWYAVRTAVINAASGR QTVDEALKDAQTNSSSAGLVPRGSGWGA<u>PLDGEYFTLQIRGRERFEMFRELNEALELKDAQAGKEPGGSRAHSSHLKSKKGQSTSRHKKLMFKTEGPDSD</u>

For mammalian-cell expression of full-length ZBTB38, the human WT ZBTB38 coding sequence was amplified from genomic DNA, digested with EcoRI and BamHI, and cloned into the corresponding sites of pEGFP-C2. The generated construct was checked by PCR and phenotypically. For the human ZBTB38 K43R mutant, a gBlock IDT fragment was digested with EcoRI and XcmI, then cloned into the corresponding sites of pEGFP-C2. The construct was controlled by DNA sequencing with the forward primer: GGCCGGACTCAGATCTCG and the reverse primer AGCTTTGAGATCGTCCAGC.

### Protein production and purification

All proteins were expressed in Rosetta (DE3) competent cells grown in 2x Yeast Extract Tryptone (2YT) medium supplemented with 50 μg/ml kanamycin (for all pET-28a and pET-24a vectors) or 100 μg/ml spectinomycin (for the plasmid encoding SAE1:SAE2). Cultures were incubated at 37 °C until the OD_600_ reached 0.8-1.0, then the expression was induced with 0.5 mM isopropylthio-β-galactoside (IPTG). After overnight expression at 18 °C, the cells were harvested and frozen at -20 °C.

Defrosted cells were resuspended in lysis buffer (500 mM NaCl, 25 mM HEPES pH 7.5, 1 mM TCEP). In the case of SAE1:SAE2, the buffer was supplemented with 0.5 mg/ml lysozyme. The cell suspension was incubated for 20 min at 37 °C, sonicated in a cold bath, and centrifuged for 30 min at 15,000 g.

The supernatant was loaded onto a 5-ml HisTrap HP column (Cytiva, cat. no. 17-5248-01) pre-equilibrated with lysis buffer. After sample loading, the column was washed with lysis buffer supplemented with 25 mM imidazole, followed by elution with an imidazole gradient (25-250 mM) over a total volume of 100 ml. The major peak fractions were pooled.

The proteins were further purified by ion exchange using a 5-ml HiTrap Q HP column (Cytiva, cat. no. 17-1154-01) for SUMO1, SAE1:SAE2, and ZBTB38^BTB^ (WT and mutants). In the cases of UBC9, RANGAP1^CTD^, and ZBTB33^BTB^ (WT and K43R), a 5-ml HiTrap SP HP column (Cytiva, cat. no. 17-1152-01) was used instead. Elution was carried out using a gradient of the NaCl concentration, from 100 to 600 mM, in 25 mM HEPES pH 7.5. The peak fractions were pooled and concentrated using Amicon with a 3-kDa (for SUMO1) or 10-kDa (for all others) cut-off. As a final purification step, proteins were buffer-exchanged either by size-exclusion chromatography using a HiLoad Superdex 75 16/600 column (Cytiva, cat. no. 28-9893-33 or 28-9893-35) equilibrated with 200 mM NaCl, 25 mM HEPES pH 7.5, and 1 mM TCEP or by repeated rounds of concentration and dilution, using an Amicon concentrator with three successive 10-fold dilution steps in the same buffer. Protein purity was confirmed using SDS-PAGE, and protein concentration was determined by the absorbance at 280 nm. Proteins were snap-frozen in liquid nitrogen and stored at -80 °C.

### Crystallisation of the WT ZBTB38

We performed crystallisation using a pure aliquot of WT ZBTB38^BTB^ protein at the final concentration of 877 μM in 25 mM HEPES pH 7.5, 150 mM NaCl and 0.5 mM TCEP. Crystallisation trials were performed at 20 °C using JCSG Plus, Morpheus I, Wizard Classic 1 + 2, and Structure 1 + 2 screens from Molecular Dimensions via the sitting-drop vapour-diffusion method using a Mosquito liquid handling instrument (TTP LabTech). Crystals were detected after a few days in Morpheus I condition 1.12: 60 mM divalent mix (MgCl_2_ and CaCl_2_), 100 mM Tris/Bicine pH 8.5 and 37.5% precipitant mix (MPD, PEG1000 and PEG3350).

### Data collection and structure determination

100-K X-ray diffraction data were collected on ID23-2 beam line at the ESRF synchrotron. The diffraction data were processed using XDS^113^ and AIMLESS^114^. The crystal structure was determined by molecular replacement using Phaser^115^ of the Phenix suite^116^. An in-house AlphaFold3^84^ prediction of a ZBTB38_1-134_ monomer was used as a successful search model. The atomic model was then refined using *phenix*.*refine* and manually improved using COOT^117^.

After final refinement rounds, R_free_ and R_work_ were calculated to be 21.52% and 23.77%, respectively. The data collection and refinement statistics are listed in Table 1. The model quality was validated by MolProbity^118^ as implemented in Phenix. Molecular graphics images were produced using Pymol^119^ or UCSF Chimera^120^.

For ConSurf analysis^85^, a representative custom multiple-sequence alignment of 84 vertebrate ZBTB4, ZBTB14, ZBTB21, ZBTB33, and ZBTB38 proteins was supplied, and conservation mapping was performed using standard settings.

### Computational protein-protein interaction screening

Protein-protein interactions were screened using a custom computational framework employing locally installed AlphaFold3^84^. Homology searches were accelerated using GPU-enabled MMseqs2, enabling efficient large-scale structural interaction screening. A single fusion protein comprising human ZBTB38 residues 1-134 repeated in tandem was used as a bait. As interaction candidates, human proteins reported to interact with full-length ZBTB38 were collected from the BioGRID database on April 29, 2026, and UBC9 was manually included in the candidate list. Sequences of all proteins were retrieved from UniProt, focussing on isoform 1 in each case. Proteins above 3,000 amino acids in length (HERC2, HUWE1) were divided into two sequences of approximately half the length each, with a short overlap. Each ZBTB38^BTB^ dimer:candidate pair was modelled only once with a single seed to efficiently use computational resources. Interaction pairs were ranked according to interface predicted template modelling (ipTM) score and the mean of two obtained interaction prediction score from aligned errors (ipSAE) scores (referred to as ipSAE_mean_).

### Analytical size exclusion chromatography (SEC)

Analytical SEC was conducted using a Superdex 75 Increase 10/300 GL column (Cytiva, cat. no. 29-1487-21) equilibrated in SEC buffer (100 mM NaCl, 25 mM HEPES, pH 7.5 & 1 mM TCEP). A 150-µL sample of protein (a ZBTB38^BTB^ variant, UBC9, or an equimolar mixture of the two) at 250 µM was loaded into a 100-µL sample loop, overfilling the loop to ensure a consistent and reproducible injection volume. Proteins were eluted isocratically at 0.5 mL/min. Elution was monitored by UV absorbance at 280 nm over 1 column volume (24 mL) and 1-mL fractions were collected. Fractions 9-15 were analysed by SDS-PAGE.

### Isothermal titration calorimetry (ITC)

ITC measurement between UBC9 and ZBTB38^BTB^ was executed at a constant temperature of 25 °C using a PEAQ-ITC instrument (Malvern instruments). Prior to the measurement, both protein solutions were equilibrated into the same buffer, 25 mM HEPES, pH 7.5, 200 mM NaCl and 1 mM TCEP, by repeated rounds of concentration and dilution in an Amicon centrifugal concentrator. The syringe contained 1 mM of ZBTB38^BTB^, whereas the cell was filled with UBC9 at 50 µM. The titration consisted of nineteen injections, each of 0.5 µL of ZBTB38^BTB^, 1 s in duration, with a 120 s interval between injections to enable the signal to return to the baseline. The stirring speed was maintained at 750 rpm. The data were analysed with the MicroCal PEAQ-ITC Analysis Software v1.41 using a single-site binding model. Experimental and fitting conditions are provided in **Figure 2D, *right***.

### SUMOylation kinetics

Reaction mixtures included protein components at the following concentrations: 20 μM SUMO1_18-97_, 0.5 μM UBC9, 0.5 μM SAE1:SAE2, and 15 μM of substrate. The reaction buffer contained 25 mM HEPES pH 7.5, 150 mM NaCl, 5% glycerol and 5 mM MgCl_2_. The reactions were performed at 37 °C and triggered by the addition of ATP at a final concentration of 5 mM. The conjugation was monitored at different time points by removing a 15-µL aliquot each time and mixing it with 15 µL of a denaturing Laemmli buffer. Proteins were separated by SDS-PAGE and visualised with ReadyBlue (Sigma Aldrich, cat. no. RSB-1L).

For quantification, gel-based assays were performed under the same conditions except for an increased amount of substrate (20 μM). Band intensities from three technical repeats were converted into kinetic curves using Image Lab software, with normalisation between the initial (0 min) and the final time points (135 min) set to 0 and 100%, respectively.

### Cell culture and transfection

RPE-1 cells were cultured in DMEM/F-12 medium (Gibco) and HEK293T cells were cultured in DMEM GlutaMAX (Gibco). Both culture media were supplemented with 10% fetal bovine serum (Gibco) and 1% penicillin-streptomycin (Gibco). Cells were maintained at 37 °C in a humidified incubator with 5% CO_2_.

For transient plasmid transfection, HEK293T cells were seeded at 5 × 10^5^ cells per well in 6-well plates and transfected immediately using Lipofectamine 2000 (Thermo Fisher Scientific). Briefly, 5 µL of Lipofectamine 2000 was diluted in 250 µL Opti-MEM (Gibco) and incubated for 5 min at room temperature. In parallel, 1 µg of plasmid DNA was diluted in 250 µL Opti-MEM. The diluted DNA and Lipofectamine solutions were combined and incubated for 10 min at room temperature to allow complex formation. 500 µL transfection mixture was then added dropwise to each well.

For siRNA transfection, RPE-1 cells were seeded at 1.5 × 10^6^ cells in T25 flask and allowed to adhere before transfection. Cells were transfected with either control or ON-TARGETplus Human ZBTB38 siRNA - SMARTpool (Horizon) using Lipofectamine RNAiMAX (Thermo Fisher Scientific). For each T25 flask, 13 µL of Lipofectamine RNAiMAX was diluted in 650 µL of Opti-MEM (Gibco) and incubated for 5–10 min at room temperature. In parallel, 3 µL of 100 µM siRNA was diluted in 650 µL of Opti-MEM. The diluted siRNA solution was then combined with the diluted Lipofectamine RNAiMAX solution and incubated for an additional 5 min at room temperature to allow complex formation. A total volume of 1.31 mL of the transfection mixture was subsequently added to each flask.

### Protein extraction and western blot

Cells were harvested 24 h after transfection and cell pellets were processed for protein extraction.

For RIPA extraction, cell pellets were lysed in RIPA buffer (Thermo Fisher Scientific) supplemented with 0.1% PMSF and a protease inhibitor cocktail (Roche). Lysates were incubated on ice for 30 min before the addition of CaCl_2_ to a final concentration of 1 mM either in the presence or absence of 10 mM N-ethylmaleimide (Sigma). To facilitate chromatin digestion, lysates were incubated for 1 min at 37 °C and treated with micrococcal nuclease (MNase; final concentration 0.00025 U/µL) for 15 min at 37 °C under agitation. The reaction was stopped by adding EDTA to a final concentration of 4 mM. Samples were subsequently sonicated using a Bioruptor (Diagenode) for 5 min (30 s ON/30 s OFF cycles, high power) to ensure complete cell lysis and chromatin fragmentation. Insoluble material was removed by centrifugation at 13,000 rpm for 10 min at 4 °C and the supernatants were collected. Protein concentrations were determined using the BCA Protein Assay Kit (Thermo Fisher Scientific).

For direct Laemmli extraction, cells were washed once with PBS before the addition of 2X Laemmli sample buffer prepared from 4X Laemmli buffer (Bio-Rad) diluted with nuclease-free water and supplemented with β-mercaptoethanol according to the manufacturer’s instructions. Cells were scraped directly into the buffer, collected, heated at 95 °C for 5 min and sonicated for 5 min (30 s ON/30 s OFF cycles, high power).

For RIPA extracts, equal amounts of protein were mixed with NuPAGE 4× LDS Sample Buffer and 10× Sample Reducing Agent (Thermo Fisher Scientific).

Before electrophoresis, all samples were denatured at 95 °C for 5 min. Proteins were separated on 4–12% NuPAGE Bis-Tris precast gels (Invitrogen) by SDS-PAGE and transferred onto nitrocellulose membranes (Millipore). Membranes were blocked in 5% (w/v) non-fat dry milk prepared in PBST (PBS containing 0.1% Tween-20) and incubated overnight at 4 °C with primary antibodies diluted in the same blocking buffer. The following primary antibodies were used: anti-ZBTB38 (HPA014865, Sigma; 1:1,000) and anti-α-tubulin (ab7291, Abcam; 1:10,000). Membranes were washed three times with PBST before incubation for 45 min at room temperature with IRDye secondary antibodies (LI-COR, 1:10,000) diluted in 5% milk in PBST. After three washes with PBST, protein signals were detected using an Odyssey Fc Imaging System (LI-COR).

## Acknowledgements

We thank beamline staff at the European Synchrotron Radiation Facility (ESRF; Grenoble, France) for access to the ID23-2 beamline and expert support. We are grateful to Jacob Seeler, Jack-Christophe Cossec, and the members of the M.J.S. team for useful discussions.

