## Supplementary Figures for "A new class of inherently efficient SUMOylation substrates"

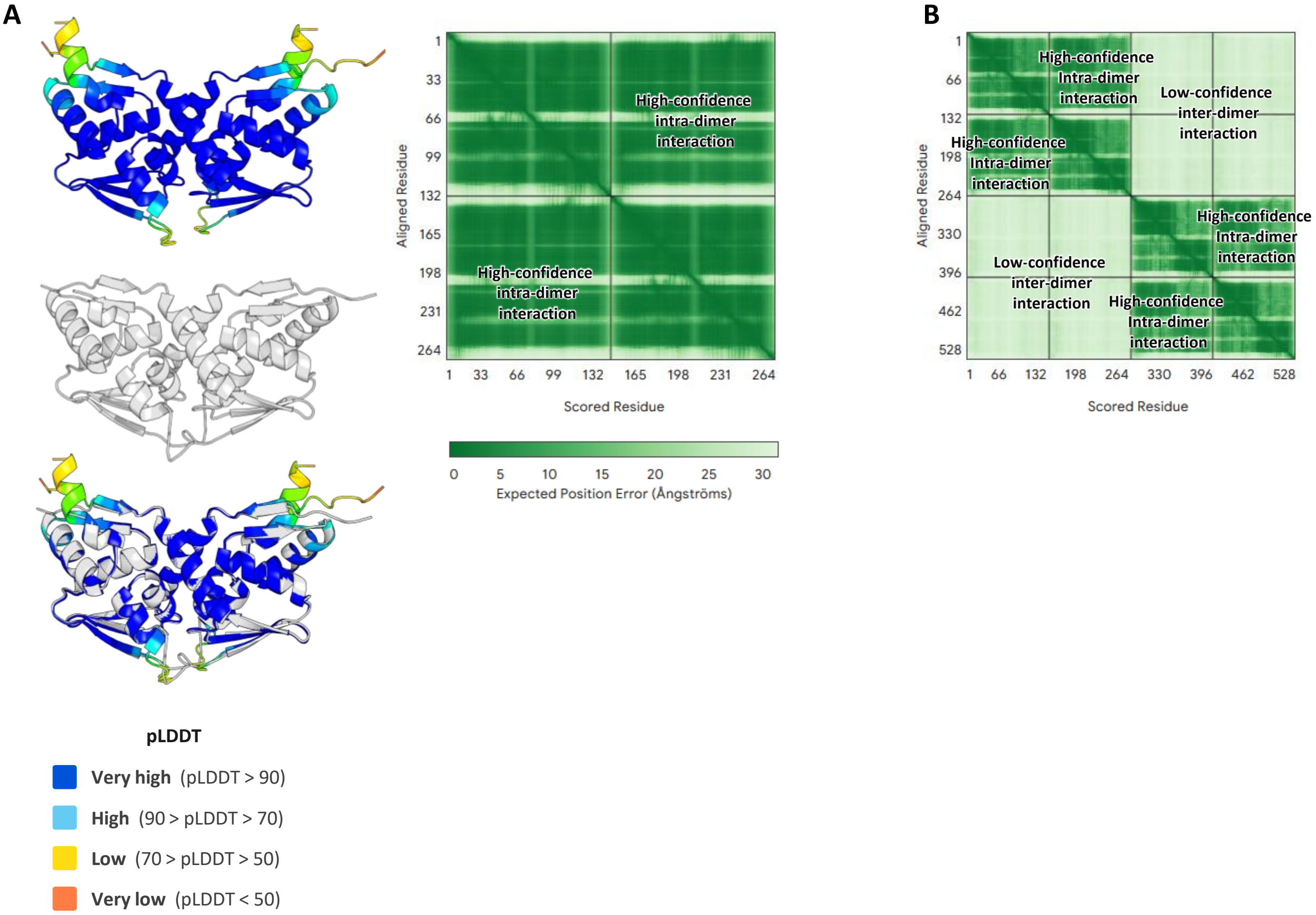


**Supplementary Figure 1. AlphaFold3 models of dimeric and putative tetrameric ZBTB38$^{BTB}$ assemblies**

(**A**) *Left*, structural alignment between the AlphaFold3-predicted (coloured) and our X-ray diffraction-derived (grey) models of the ZBTB38$^{BTB}$ dimer, shown first individually, and then, below, super-imposed. The AlphaFold3 model is coloured according to the per-residue predicted local distance difference test (pLDDT) score, using the scale indicated underneath. *Right*, predicted aligned error (PAE) plot for the AlphaFold3 dimer prediction with position error represented using the colour scale provided below. The off-diagonal squares indicative of inter-protomer (intra-dimer) interactions are labelled.

(**B**) PAE plot for the AlphaFold3 prediction of a putative tetrameric assembly, coloured like in panel **A**. The off-diagonal squares representing intra- and inter-dimer interactions are labelled.

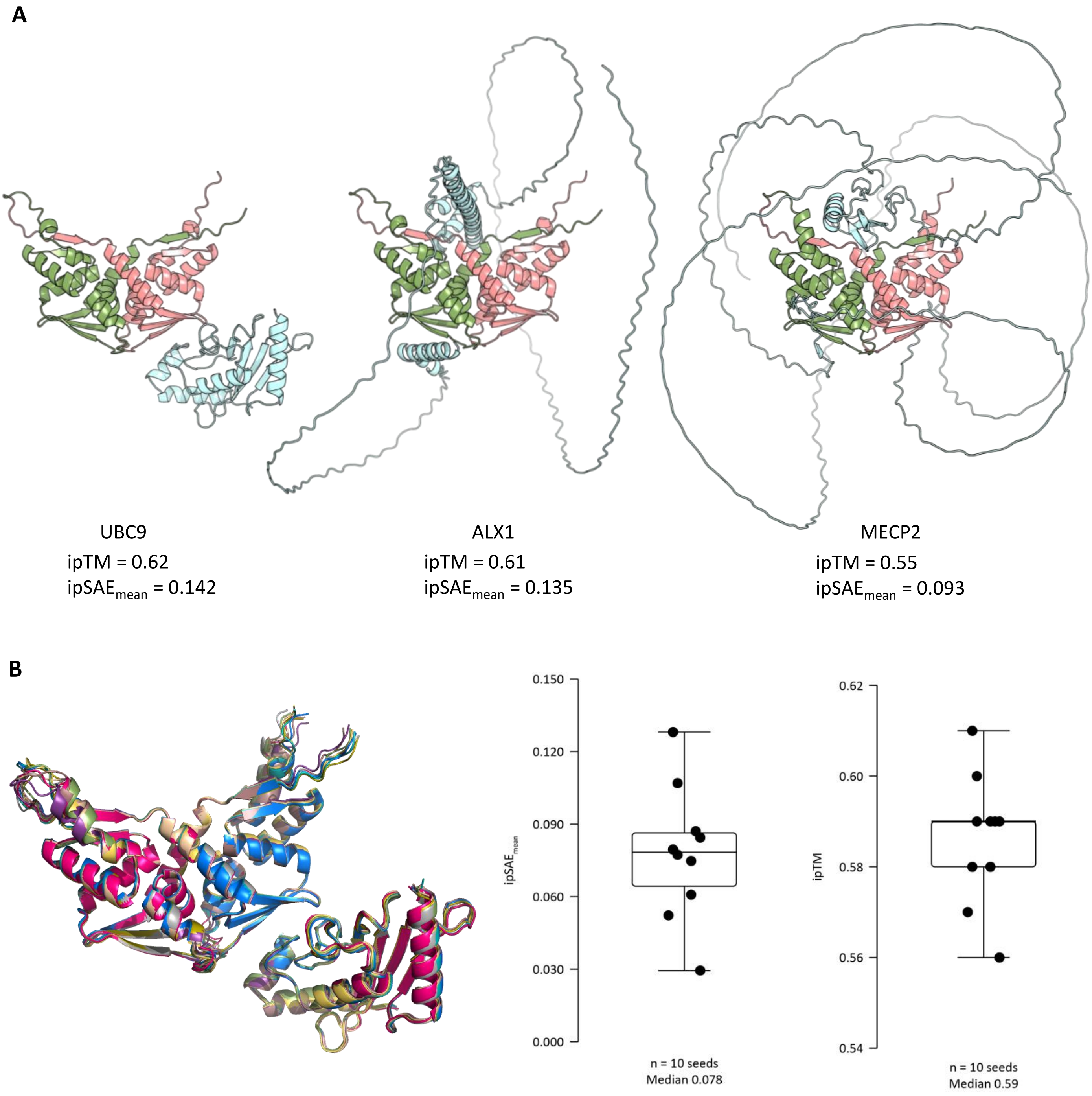


**Supplementary Figure 2. Top-scoring AlphaFold3-predicted ZBTB38$^{\text{BTB}}$-dimer interactions**

(A) Cartoon representations of three top-scoring models from the AlphaFold3 ZBTB38$^{\text{BTB}}$-dimer interaction screen in Figure 1D. The ZBTB38$^{\text{BTB}}$ dimer was modelled as a single chain composed of two fused protomers (coloured green and pink) arranged in tandem. The indicated interactors are shown in blue. Interactor names and predicted confidence metrics – ipTM and $\text{ipSAE}_{\text{mean}}$ – are provided underneath.

*(B)* *Left*, ten AlphaFold3 models of the ZBTB38$^{\text{BTB}}$:UBC9 complex generated with different random seeds (each coloured differently), superposed on one another. *Right*, box plots showing mean interaction prediction score from aligned errors ($\text{ipSAE}_{\text{mean}}$) and interface predicted template modelling (ipTM) scores calculated for these models. Points show scores for individual models. The boxes span the interquartile range (25th–75th percentiles) with the median marked inside (note that for ipTM, the median nearly coincides with the upper box rim). Median values are provided underneath. Whiskers indicate data extremes.

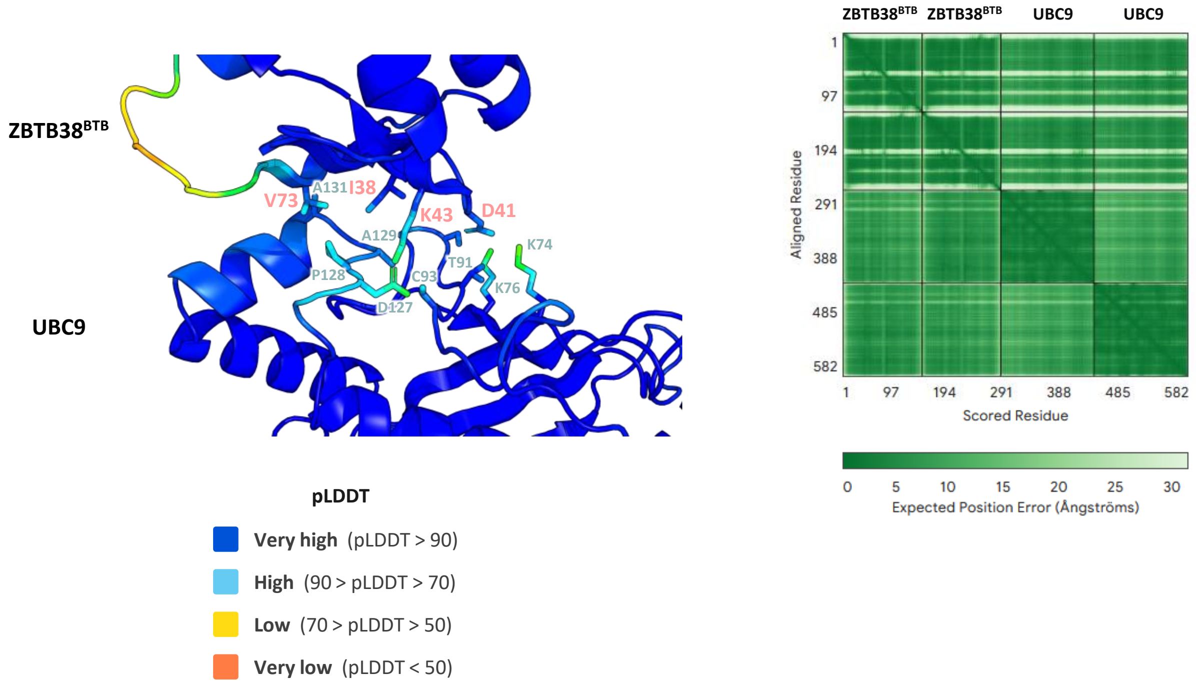


**Supplementary Figure 3. Additional predicted confidence metrics for the ZBTB38$^{BTB}$:UBC9 model**

*Left*, cartoon representations of the AlphaFold3-modelled ZBTB38$^{BTB}$:UBC9 interface coloured according to the per-residue predicted local distance difference test (pLDDT) score, using the scale provided below. Selected interface side chains are shown as sticks and labelled with pink (ZBTB38$^{BTB}$) or blue (UBC9) labels.

*Right*, the predicted aligned error (PAE) plot for the AlphaFold3-predicted ZBTB38$^{BTB}$:UBC9 complex in the 2:2 stoichiometry, with position error represented using the below-provided colour scale. The sequence fragments corresponding to the individual ZBTB38$^{BTB}$ and UBC9 chains are labelled above the PAE plot.
